# Highly selective transgene expression through double-floxed inverted orientation system by using a unilateral spacer sequence

**DOI:** 10.1101/2022.04.11.487972

**Authors:** Natsuki Matsushita, Shigeki Kato, Kayo Nishizawa, Masateru Sugawara, Kosei Takeuchi, Yoshiki Miyasaka, Tomoji Mashimo, Kazuto Kobayashi

## Abstract

The double-floxed inverted orientation (DIO) system with adeno-associated viral (AAV) vector provides a beneficial approach to express transgenes in specific cell populations having Cre recombinase. A significant issue with this system is the protection against non-specific expression of transgenes in tissues after vector injection. We here show that Cre-independent recombination in AAV genome carrying the DIO sequence occurs during the production of viral vectors in packaging cells, which results in transgene expression in off-target populations. Introduction of a relatively longer nucleotide sequence between two recognition sites at the unilateral side of the transgene cassette, termed a unilateral spacer sequence (USS), is useful to suppress recombination during the vector production, leading to the protection of non-specific transgene expression with enhanced gene expression selectivity. Our DIO/USS system offers a powerful strategy for highly specific Cre-dependent transgene expression, aiming at various applications for structural and functional analyses of target cell populations.

## INTRODUCTION

Cre-*lox*P-mediated recombination is a system for site-specific recombination derived from bacteriophage P1 (Sternberg et al., 1981; Abremski et. al., 1983) and has been used for the research of a variety of experimental biology (Sauer, 1987; Sauer and Henderson, 1988; Gu et al., 1993). In this recombination system, Cre recombinase catalyzes site-specific recombination between two sites of *lox*P, which is a 34-base pair (bp) DNA sequence containing an 8-bp asymmetrical spacer flanked by two 13-bp inverted repeats (Lee and Saito, 1998; Hoess et al., 1999; Voziyanov et al., 1999). An application of the Cre-*lox*P system has been developed to express genes of interest in specific cell populations, termed the double-floxed inverted orientation or open reading frame (DIO) system or the flip-excision switch system (Saunders et al., 2012; Schnütgen et al., 2003; Sohal et al., 2009) (see Figure S1). In this system, a gene cassette in an inverted orientation is flanked by a pair of double recognition sites, which consist of *lox*P and *lox*2272 sites with a ∼50-bp spacer, in the opposite direction with an alternate order at both ends of the cassette. These recognition sites are different in two nucleotides in the 8-bp asymmetrical spacer, and are exclusively recombined by Cre recombinase. The recombinase first catalyzes reversion of the gene cassette through recombination between either of the double recognition sites, and generates a pair of either the *lox*P or *lox*2272 site in the same orientation with an intervening sequence containing another recognition site at either end of the cassette, leaving a single recognition site corresponding to another one at the opposite end. In the second reaction, the recombinase catalyzes deletion of the intervening sequence via recombination between the pair of recognition sites, and produces the reversed gene cassette with different recognition sites at both ends, preventing re-inversion of the cassette.

Adeno-associated viral (AAV) vectors are widely used to apply the DIO system, in particular in the research field of neuroscience, aiming at transgene expression in specific cell populations dependent on Cre recombinase by using transgenic animals expressing the recombinase under the control of cell type-specific gene promoters (Kozorovitskiy et al., 2012; Miyamichi et al., 2013; Nonomura et al., 2018; Srinivasan et al., 2016; O’Connor et al., 2013) or viral vectors expressing the recombinase in specific neural circuits (Aelvoet et al., 2015; Arguello et al., 2017; Kato et al., 2018; Tschida et al., 2019; Woods et al., 2020). However, there is evidence for non-specific transgene expression in off-target cell populations after treatment of AAV vectors carrying the DIO system, which appears to arise at least from infrequent transcription from the inverted transgene in viral genome and recombination events through short homologous sequence within recognition sites during transfer plasmid amplification in bacterial cells (Fischer et al., 2019). To resolve these issues, AAV vectors with mutated recognition sites with the decreased homology were developed to suppress bacterial recombination events, combined with the in-frame initiation codon located upstream of the recognition sites at the 5’-end of the transgene, in which Cre-mediated recombination deletes a gene cassette containing polyadenylation signals flanked by the two recognition sites in the same orientation (Fischer et al., 2019). Another possibility to explain non-specific transgene expression in the DIO system may be recombination events independent of Cre recombinase accompanied by reversion of the inverted transgene during the production of AAV vectors in packaging cells. The recombination in viral genome during AAV production in the cells has not yet been fully investigated.

Protection of non-specific transgene expression in off-targets is an important issue for application of the DIO system to achieve highly selective gene expression dependent on Cre recombinase. In the present study, we found that recombination events indeed occur between a pair of nucleotide sequences within recognition sites during the production of AAV vectors with the DIO sequence in cultured cells, although the recombination during the amplification of transfer plasmid in bacterial cells is not detectable. We then found that the introduction of a relatively longer nucleotide sequence (750 bp) between the *lox*P and *lox*2272 sites at the unilateral side of the inverted transgene cassette, termed a unilateral spacer sequence (USS), is useful to suppress the recombination in viral genome during vector production, which results in the protection of non-specific transgene expression with enhanced selectivity of Cre-dependent gene expression in brain tissues. A DIO system that includes a USS is applicable for the highly selective expression of several types of transgenes in target cell populations.

## RESULTS

### Cre-independent recombination during AAV vector production results in non-specific expression of transgene in tissues

We used an AAV serotype 2 vector encoding the gene cassette for green fluorescent protein (GFP) in the inverted orientation, which is flanked by a pair of double recognition sites between the cytomegalovirus early enhancer/chicken *β*-actin (CAG) promoter (Niwa et al., 1991) and woodchuck hepatitis virus posttranscriptional regulatory element (WPRE) connected to human growth factor (hGH) gene polyadenylation signal, termed AAV-DIO-GFP vector (see Figure 1A for the genome structure and Figure S2 for the nucleotide sequence). Prior to the production of AAV vector, we checked the presence of recombinants with the reversion of inverted GFP gene cassette in the transfer gene plasmid pAAV-DIO-GFP preparation with polymerase chain reaction (PCR), and this type of recombinant was not detectable in our plasmid preparation (see Figure S3). HEK293T cells were transfected with the plasmids encoding the transfer gene, replication/encapsulation (RC) genes, and adeno-helper genes, and vector particles were collected from the supernatant of culture medium. To test whether vector particles contain recombinant forms (RFs) carrying viral genome with the reversed GFP gene cassette, we measured the total and recombinant vector titers by using quantitative PCR with the following primer sets and probes (Figure 1A, B). The quantitative PCR with the primer set FP-W/RP-W1, which amplifies a part of the WPRE sequence, and the primer set FP-G1/RP-W2, which amplifies a part containing the 3’-region of a reversed GFP sequence, provided a total vector titer of 3.16 `±` 0.43 × 10^12^ genome copies/mL and a recombinant vector titer of 3.65 `±` 0.48 × 10^10^ genome copies/mL (n = 4 samples), respectively. The frequency of recombination, defined as the percentage of the recombinant vector titer divided by the total vector titer, was 1.16 `±` 0.02% (n = 4 samples).

**Figure 1.**
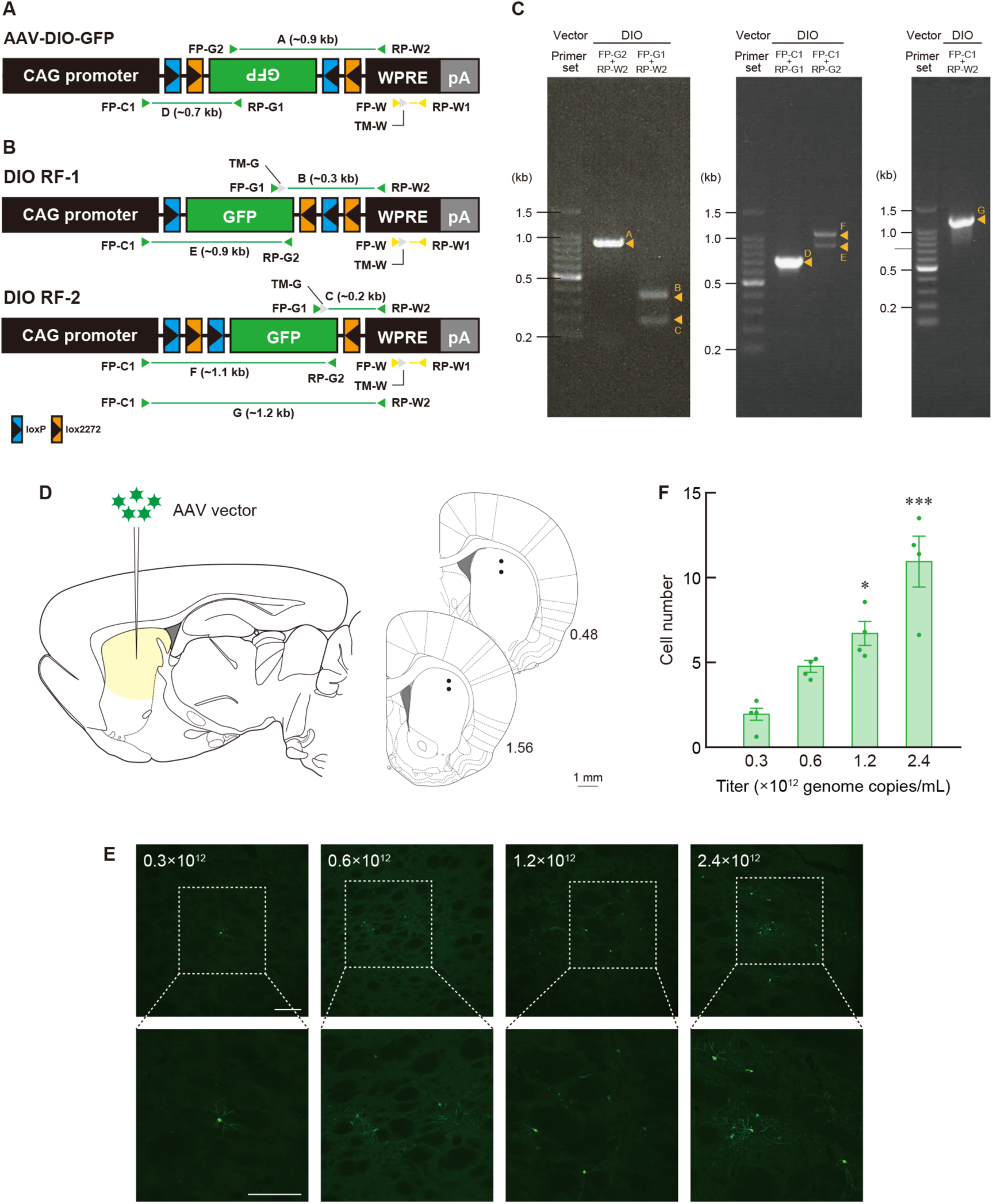
Cre-independent recombination during vector production and non-specific transgene expression after vector injection. (A) Genome structure of AAV-DIO-GFP vector. The GFP gene cassette in the inverted orientation is flanked by two double recognition sites and located between the CAG promoter and WPRE sequence connected to hGH polyadenylation signal (pA). (B) Genome structures of RFs contained in AAV-DIO-GFP vector particles. Arrowheads indicate the position and direction of forward primers (FP), reverse primers (RP), and TaqMan probes (TM) for PCR and quantitative PCR amplification. DNA fragments with their sizes obtained from PCR amplification are indicated. (C) Analysis of amplified DNA fragments. PCR amplification was carried out with viral genome of 1.0 × 10^6^ genome copies as a template by using the indicated primer sets, and PCR products were subjected to 1% agarose gel electrophoresis. Fragments B, C, E, and F were used for detrmination of nucleotide sequences (Figure S4A, B). (D) Schematic illustration for intracranial injection of AAV vector. Anteroposterior coordinates (mm) from bregma in rats are shown, and dots indicate the injection sites in the striatum. (E) GFP immunohistochemistry with striatal sections. Wild-type Long Evans rats were unilaterally given intracranial injection of the vector with different genome titers (0.3 × 10^12^ to 2.4 × 10^12^ genome copies/mL) into the striatum (0.1 μL/site × 4 sites), and sections through the striatum were immunostained with anti-GFP antibody. Representative images of GFP immunostaining are indicated for each vector titer. Lower images show magnified views of the squares in the upper images. (F) Titer-dependent increase in the number of GFP^+^ cells. Data are presented as mean ± SEM (n = 4 animals). Individual data are overlaid. \**P* < 0.05 and \*\*\**P* < 0.001 compared with the lowest titer (Bonferroni test). Scale bars: 1 mm (D); and 200 μm (E).

To characterize the structures of viral genomes of the RFs in detail, we carried out PCR amplification of vector particles with different primer sets (Figure 1A, B) and detected PCR products using 1% agarose gel electrophoresis (Figure 1C). The FP-G2/RP-W2 primer set generated a DNA band of ∼0.9 kilobases (kb), which corresponds to fragment A in the original viral genome. The primer set FP-G1/RP-W2 provided two kinds of DNA band, ∼0.3 kb and ∼0.2 kb, which respectively correspond to fragments B and C in the RF genomes with the reversion of the GFP gene cassette. The FP-C1/RP-G1 primer set gave a DNA band of ∼0.7 kb, which corresponds to the 5’-part of the original genome (fragment D), and the FP-C1/RP-G2 primer set produced two DNA bands, ∼0.9 kb and ∼1.1 kb, equivalent to fragments E and F, respectively, in the RFs. Nucleotide sequence analysis of fragments B and E indicated the primary structure of RF-1, which was composed of the reversal of the inverted GFP sequence flanked by a single *lox*P site at the 5’-end and *lox*2272-*lox*P-*lox*2272 sites at the 3’-end (Figure S4A), and the analysis of fragments C and F showed the structure of RF-2, which contained the reversed GFP sequence with *lox*P-*lox*2272-*lox*P sites at the 5’-end and a single *lox*2272 site at the 3’-end (Figure S4B). The structures of RF-1 and RF-2 coincided with those of the recombinants formed through the first recombination in the DIO system, suggesting that these RFs seem to be produced, in the absence of Cre recombinase, through recombination events between a pair of nucleotide sequences within the *lox*P or *lox*2272 sites during AAV vector production in cultured cells (see Figure S5 for possible mechanisms to explain the recombination events). In addition, PCR amplification with the FP-C1/RP-W2 primer set, which amplifies a DNA fragment from the 3’-portion of CAG promoter to the 5’-portion of WPRE, displayed only a single band of ∼1.2 kb (fragment G), and did not detect any smaller bands corresponding to DNA fragments deduced from recombination (deletion) between a pair of recognition sites in the same orientation (Figure 1C), suggesting that the recombination between the two recognition sites rarely occurs during vector production.

Next, to examine whether the RFs in AAV-DIO-GFP vector particles cause non-specific transgene expression in the brain tissues of wild-type rats, we performed an intracranial injection of the vector with different total titers (0.3 × 10^12^ to 2.4 × 10^12^ genome copies/mL) into the striatum (1.0 μL/site × 4 sites) of Long Evans rats (see Figure 1D for the injection coordinates). Sections through the striatum were stained by immunohistochemistry with anti-GFP antibody. Representative images of immunostaining are shown in Figure 1E. The number of GFP^+^ cells in regions of interest (ROIs; 1.0 × 1.0 mm) in the striatum was counted and the average cell number was calculated (see Figure S6 for ROIs). The cell number was elevated gradually, along with the increasing vector titers (Figure 1F; one-way ANOVA, *F*_3,12_ = 19.621, *P* < 0.001). The number at the higher titers of 1.2 × 10^12^ and 2.4 × 10^12^ genome copies/mL (6.72 ± 0.70 and 10.97 ± 1.48 cells, respectively, n = 4 animals) indicated significant increases as compared to that at the lowest titer of 0.3 × 10^12^ genome copies/mL (1.94 ± 0.42 cells, n = 4 animals) (Bonferroni test; *P* < 0.05 for 1.2 × 10^12^ genome copies/mL, *P* < 0.001 for 2.4 × 10^12^ genome copies/mL). These data show that the RFs in vector particles result in titer-dependent, non-specific transgene expression after AAV vector treatment of the wild-type brains.

### Suppression of Cre-independent recombination protects non-specific transgene expression through the DIO system by using the USS

To perform the selective transgene expression based on the DIO system, we need to suppress Cre-independent recombination in viral genome during vector production in cultured cells and non-specific transgene expression in tissues after vector injection. In our preliminary experiments, we generated a modified AAV vector carrying the DIO system, in which the gene encoding TurboFP635 (Shcherbo et al., 2007) was introduced between the *lox*P and *lox*2272 sites at the 5’-end of the inverted GFP cassette to develop a strategy to distinguish Cre^+^ and Cre^-^populations in cells transduced by the AAV vector. In this strategy, we aimed to label Cre^-^ cells with TurboFP635 expression and to induce GFP expression in Cre^+^ cells through Cre-dependent recombination. During the progress of experiments, we observed that the introduction of the TurboFP635 gene between double recognition sites suppressed non-specific GFP expression following AAV vector injection into the brains of the wild-type rats. Based on these observations, we hypothesized that spacing between double recognition sites with a relatively longer sequence may be able to prevent recombination events in AAV genome with the DIO sequence in cultured cells. To test this possibility, we introduced the nucleotide sequence of 750 bp containing the TurboFP635 gene between the *lox*P and *lox*2272 sites either at the 5’- or 3’-end as a USS, resulting in AAV-DIO/5’USS-GFP or AAV-DIO/3’USS-GFP vector (see Figure 2A for the structure and Figure S7A, B for the nucleotide sequences). These AAV vectors were produced and vector particles were collected as described above. Total and recombinant vector titers were measured by using the quantitative PCR similarly to the case of AAV-DIO-GFP vector (Figure 2A, B). The frequency of recombination showed a significant difference among the three vectors (Table 1; one-way ANOVA, *F*_2,9_ = 92.177, *P* < 0.001), and the frequency was markedly reduced in AAV-DIO/5’USS-GFP (0.15 ± 0.04%, n = 4 samples) and AAV-DIO/3’USS-GFP (0.02 ± 0.01%, n = 4 samples) vectors as compared to AAV-DIO-GFP vector (1.22 ± 0.11%, n = 4 samples) (Bonferroni test, *P* < 0.001 vs AAV-DIO-GFP). In particular, the extent of reduction was greater in AAV-DIO/3’USS-GFP vector than AAV-DIO/5’USS-GFP vector.

**Figure 2.**
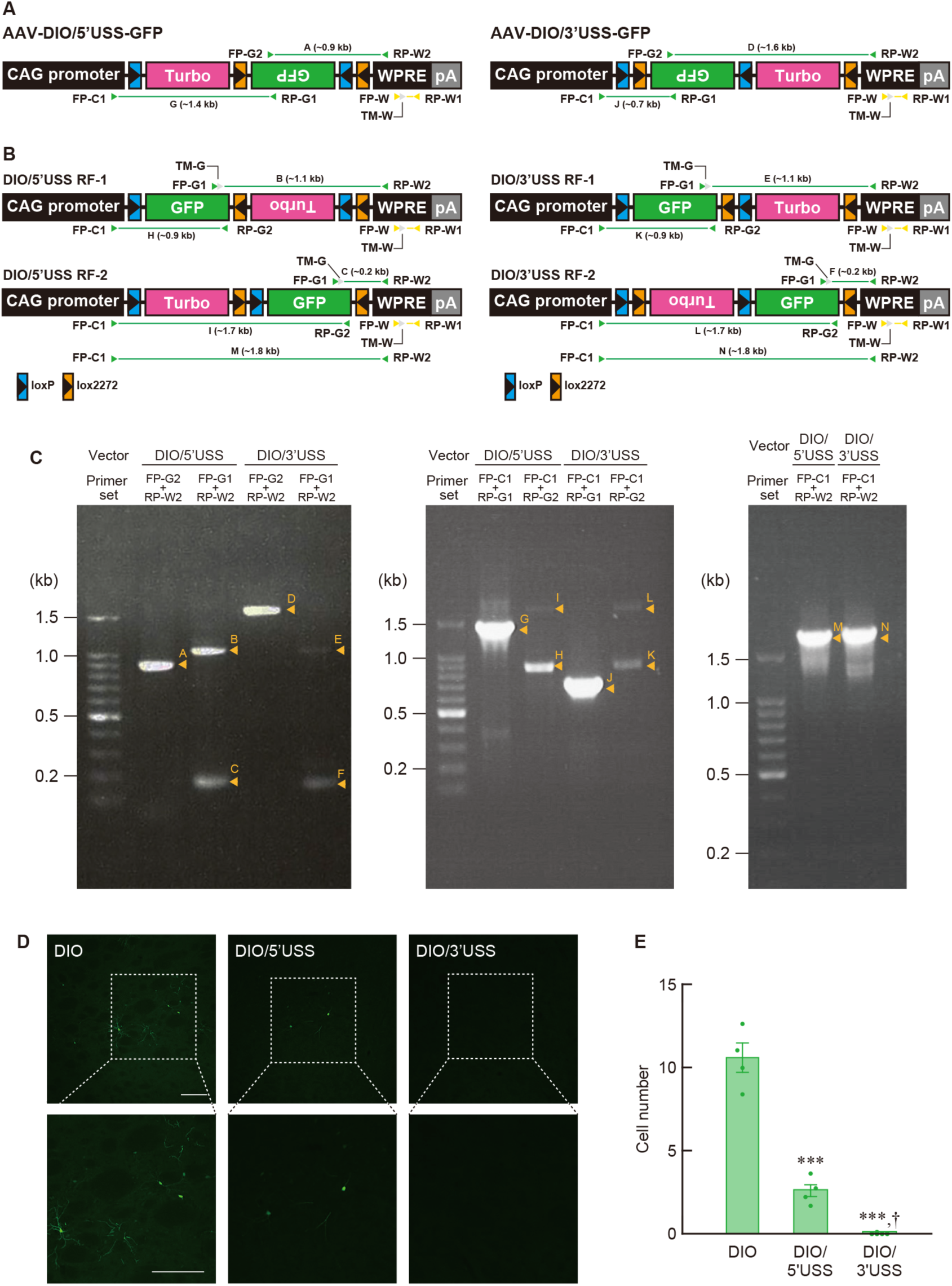
Suppression of Cre-independent, non-specific transgene expression by the use of the USS. (A) Genome structures of AAV-DIO/5’USS-GFP and AAV-DIO/3’USS-GFP vectors. The TurboFP635 sequence as the USS was introduced between the *lox*P and *lox*2272 sites either at the 5’- or 3’-end of the inverted GFP gene cassette in AAV-DIO-GFP vector, resulting in AAV-DIO/5’USS- GFP and AAV-DIO/3’USS-GFP vectors. pA: hGH polyadenylation signal. (B) Genome structures of RFs in AAV-DIO/5’USS-GFP and AAV-DIO/3’USS-GFP vector particles. Arrowheads indicate the position and direction of forward primers (FP), reverse primers (RP), and TaqMan probes (TM) for PCR and quantitative PCR amplification. DNA fragments with their sizes obtained from PCR amplification are presented. (C) Analysis of amplified DNA fragments. PCR amplification was carried out with viral genome of 1.0 × 10^6^ genome copies as a template by using the indicated primer sets, and PCR products were subjected to 1% agarose gel electrophoresis. Fragments B, C, E, F, H, I, K, and L were used for nucleotide sequence determination (Figure S8A–D). (D) GFP immunohistochemistry with striatal sections. Long Evans rats received the intracranial injection of three kinds of vectors with titer of 2.4 × 10^12^ genome copies/mL into the striatum (0.1 μL/site × 4 sites), and striatal sections were used for GFP immunostaining. Representative images of GFP staining are shown for each vector. Lower images are magnified views of the squares in the upper images. (E) Introduction of the USS into the vector markedly reduced the number of GFP^+^ cells. Data are presented as mean ± SEM (n = 4 animals). Individual data are overlaid. \*\*\**P* < 0.001 compared with AAV-DIO-GFP vector, ^†^*P* < 0.05 compared with AAV-DIO/5’USS-GFP vector (Bonferroni test). Scale bars: 200 μm (D).

**Table 1.** Titration of AAV vectors.

| Vector | Total vector titer<br>(genome copies/mL) | Recombinant vector titer<br>(genome copies/mL) | Recombination<br>frequency (%) |
| --- | --- | --- | --- |
| AAV-DIO-GFP | $2.76 \pm 0.34 \times 10^{12}$ | $3.29 \pm 0.25 \times 10^{10}$ | $1.22 \pm 0.11^{**}$ |
| AAV-DIO/5'USS-GFP | $5.79 \pm 1.05 \times 10^{12}$ | $8.71 \pm 3.06 \times 10^9$ | $0.15 \pm 0.04^{***}$ |
| AAV-DIO/3'USS-GFP | $2.20 \pm 0.70 \times 10^{13}$ | $2.65 \pm 0.75 \times 10^9$ | $0.02 \pm 0.01^{***}$ |

Total and recombinant vector titers in vector preparations were measured by using quantitative PCR with different primer sets and probes. Recombination frequency was defined as the percentage of the recombinant vector titer divided by the total vector titer.
n = 4 samples. \*\*\* $P < 0.001$ compared with AAV-DIO-GFP vector (Bonferroni test).

We characterized the viral genome structures of the RFs in AAV-DIO/5’USS-GFP and AAV-DIO/3’USS-GFP vector particles by using PCR amplification with the same primer sets as those used for AAV-DIO-GFP vector (Figure 2A, B). PCR products were detected by 1% agarose gel electrophoresis (Figure 2C). First, the structures of the RFs in AAV-DIO/5’USS-GFP particles were analyzed. The FP-G2/RP-W2 primer set gave a DNA band of ∼0.9 kb, which corresponds to fragment A in the original viral genome, and the FP-G1/RP-W2 primer set generated two DNA bands of ∼1.1 kb and ∼0.2 kb, corresponding to fragments B and C, respectively, in the RF genomes. The FP-C1/RP-G1 primer set gave a DNA band of ∼1.4 kb, which corresponds to fragment G in the 5’-region of the original genome, and the FP-C1/RP-G2 primer set produced two DNA bands of ∼0.9 kb and ∼1.7 kb equivalent to fragments H and I, respectively, in the 5’-region of the RFs. Nucleotide sequence determination of fragments B and H indicated the primary structure of RF-1, which consisted of the reversion of the TurboFP635-*lox2272*-inverted GFP sequence flanked by a single *lox*P site at the 5’-end and *lox*P-*lox*2272 sites at the 3’-end (Figure S8A), and the analysis of fragments C and I had a similar structure to that of RF-2, which contained the TurboFP635 sequence flanked by single *lox*P and *lox*2272 sites at the 5’ and 3’-ends respectively, connecting to the reversed GFP sequence flanked by single *lox*P and *lox*2272 sites at the respective 5’- and 3’-ends (Figure S8B). Secondly, the RF structures in the AAV-DIO/3’USS-GFP particles were analyzed. The FP-G2/RP-W2 primer set gave a DNA band of ∼1.6 kb (fragment D in the original genome), and the FP-G1/RP-W2 primer set generated two DNA bands of ∼1.1 kb and ∼0.2 kb (fragments E and F in the RFs, respectively). The FP-C1/RP-G1 primer set gave a DNA band of ∼0.7 kb (fragment J in the 5’-region of the original genome), and the FP-C1/RP-G2 primer set produced two DNA bands of ∼0.9 kb and ∼1.7 kb (fragments K and L in the 5’-region of the RFs, respectively). Nucleotide sequencing of fragments E and K showed the primary structure of RF-1, which consisted of the reversed GFP sequence flanked by single *lox*P and *lox*2272 sites at the 5’- and 3’-ends, respectively, linking to the TurboFP635 sequence flanked by single *lox*P and *lox*2272 sites at the respective 5’- and 3’-ends (Figure S8C), and the sequencing of fragments F and L displayed the same structure as that of RF-2, which included the inverted TurboFP635-*lox*2272-GFP sequence flanked by *lox*2272-*lox*P sites at the 5’-end and by a single *lox*2272 site at the 3’-end (Figure S8D). In addition, PCR amplification with the FP-C1/RP-W2 primer set produced only a single major band of ∼1.8 kb (fragments M and N for AAV-DIO/5’USS-GFP and AAV-DIO/ 3’USS-GFP, respectively), showing no smaller bands equivalent to DNA fragments deduced from recombination between a pair of recognition sites in the same orientation (Figure 2C). These results suggest that RF-1 and RF-2 in AAV-DIO/5’USS-GFP or AAV-DIO/3’USS-GFP particles, similar to the case of AAV-DIO-GFP vector, are generated via recombination events between nucleotide sequences within either of the double recognition sites during the production of these two vectors, and that the deletion between recognition sites in the same direction does not efficiently happen during vector production (see Figure S9 for possible mechanisms to explain the recombination events). However, the frequency of recombination in AAV-DIO/5’USS-GFP and AAV-DIO/ 3’USS-GFP particles was largely declined as compared to that in AAV-DIO-GFP vector (Table 1), suggesting that the presence of the USS between the *lox*P and *lox*2272 sites in the DIO system interferes with Cre-independent recombination events during vector production.

We then investigated whether the declined recombination frequency by the use of USS indeed protects non-specific expression of transgenes in the wild-type brains. AAV-DIO-GFP, AAV-DIO/5’USS-GFP, and AAV-DIO/3’USS-GFP vectors with equivalent total titers (2.4 × 10^12^ genome copies/mL) were injected into the striatum (1.0 μL/site × 4 sites) of Long Evans rats, and striatal sections were stained by GFP immunohistochemistry. Representative images of immunostaining are shown in Figure 2D. The number of GFP^+^ cells showed a significant difference among three kinds of vectors (Figure 2E; one-way ANOVA, *F*_2,9_ = 98.965, *P* < 0.001). The cell number in AAV-DIO/5’USS-GFP (2.63 ± 0.41 cells, n = 4 animals) and AAV-DIO/3’USS-GFP (0.03 ± 0.03 cells, n = 4 animals) was significantly reduced compared to that in AAV-DIO-GFP vector (10.59 ± 0.87 cells, n = 4 animals) (Bonferroni test, *P* < 0.001), and the number of AAV-DIO/3’USS-GFP was further lower than that in AAV-DIO/5’USS-GFP (Bonferroni test, *P* < 0.05). These data demonstrate that the use of USS in the DIO system has a suppressive effect on non-specific transgene expression in wild-type brains, and that 3’USS is more efficient than 5’USS to protect non-specific gene expression.

### Highly selective Cre-dependent transgene expression by the DIO/USS system

To ascertain whether the selectivity of Cre-dependent transgene expression is improved by the DIO system with the USS, we examined the efficacy of USS introduction on the selectivity of Cre-dependent transgene expression in the brain tissues. In this experiment, we compared the efficacy of AAV-DIO/3’USS-GFP vector to AAV-DIO-GFP vector, because this vector showed a greater effect on the protection of non-specific transgene expression as compared with AAV-DIO/5’USS-GFP vector. AAV-DIO-GFP and AAV-DIO/3’USS-GFP vectors (2.4 × 10^12^ genome copies/mL) were injected into the striatum (1.0 μL/site × 4 sites) of knock-in transgenic rats expressing Cre under the control of the gene encoding parvalbumin (PV) (Yoshimi et al., 2020). Double immunohistochemistry for PV and Cre indicated that Cre transgene was expressed in all PV-containing neurons in the striatum of PV-Cre knock-in rats (see Figure S10). Sections through the striatum were used for double immunohistochemistry for PV and GFP, and typical images of immunostaining are shown in Figure 3A. The percentages of the number of PV^+^ + GFP^+^ cells divided by that of PV^+^ cells were 93.53 ± 0.53% for AAV-DIO-GFP vector (n = 4 animals) and 95.00 ± 1.24% for AAV-DIO/3’USS-GFP vector (n = 4 animals) (Figure 3B; Student’s *t* test, *t*_6_ = 1.092, *P* = 0.317), indicating the efficient expression of GFP in PV-containing neurons in both vector-injected rats. In contrast, the percentage of the number of PV^+^ + GFP^+^ cells divided by that of GFP^+^ cells was significantly increased in AAV-DIO/3’USS-GFP vector (89.97 ± 0.69%) compared to that in AAV-DIO-GFP vector (62.34 ± 1.24%), whereas the percentage of the number of PV^-^ + GFP^+^ cells divided by that of GFP^+^ cells was markedly decreased in AAV-DIO/3’USS-GFP vector (10.03 ± 0.69%) compared to AAV-DIO-GFP vector (37.66 ± 1.24%) (Figure 3C; Student’s *t* test, *t*_6_ = 19.435, *P* < 0.001), indicating the improved selectivity of Cre-dependent transgene expression by the DIO system with 3’USS.

**Figure 3.**
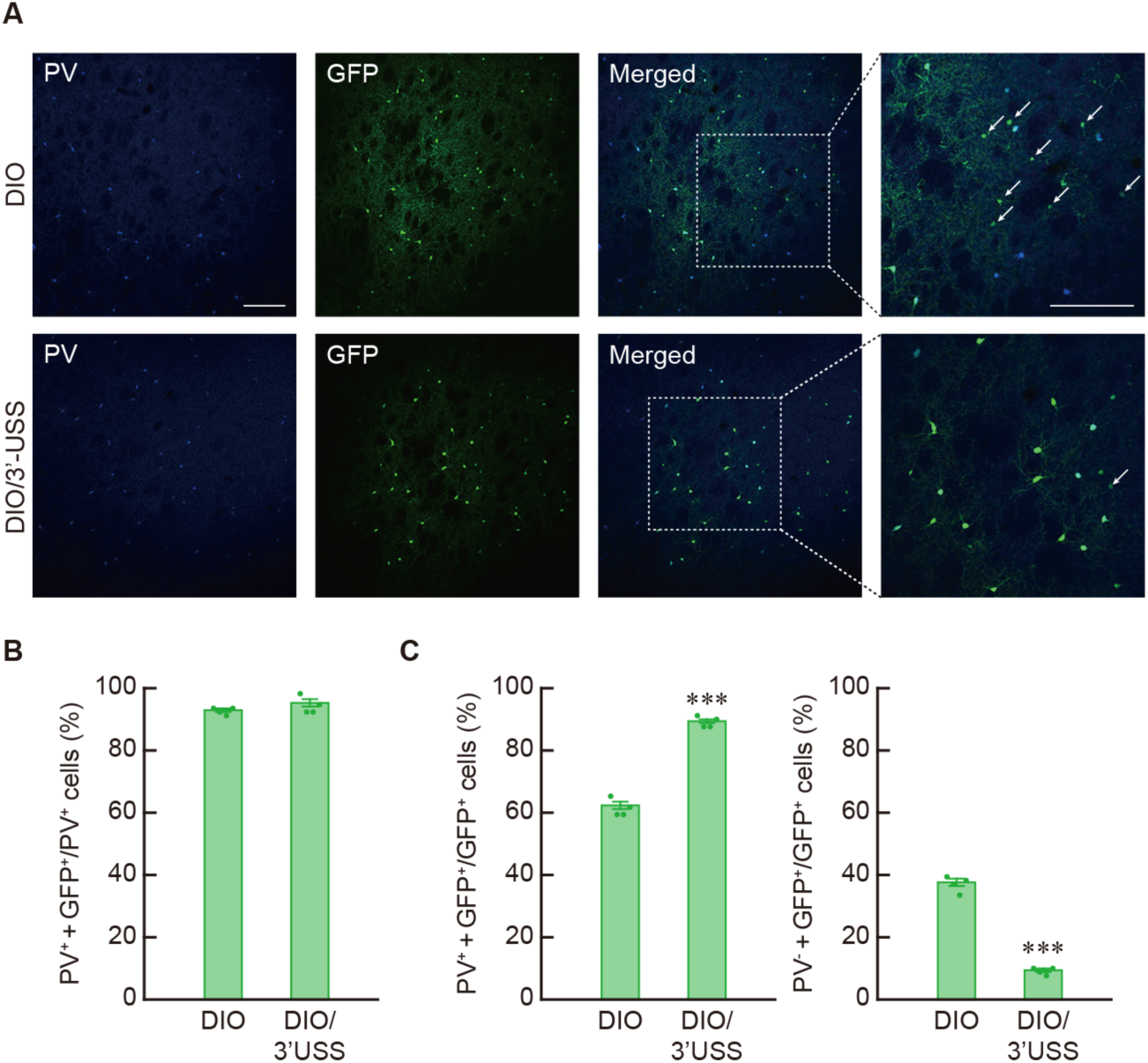
Improved selectivity of Cre-dependent transgene expression through the DIO/USS system. (A) Double immunohistochemistry for PV and GFP with striatal sections. PV-Cre knock-in rats were given unilaterally the intracranial injection of AAV-DIO-GFP or AAV-DIO/3’USS-GFP vector (2.4 × 10^12^ genome copies/mL) into the striatum (0.1 μL/site × 4 sites), and sections through the striatum were used for double immunohistochemistry with anti-PV and anti-GFP antibodies. Typical images of double immunostaining are presented for each vector. Right photos are magnified views of the squares in the merged photos. Arrows indicate PV^-^/GFP^+^ cells. (B) High frequency of Cre-dependent expression by the DIO/USS system. The percentage of the number of PV^+^/GFP^+^ cells to that of PV^+^ cells was calculated. (C) Improved selectivity of Cre-dependent expression by the use of the USS. The percentage of the number of PV^+^/GFP^+^ cells to that of GFP^+^ cells and the number of PV^-^/GFP^+^ cells compared to that of GFP^+^ cells were calculated. Data are presented as mean ± SEM (n = 4 animals). Individual data are overlaid. \*\*\**P* < 0.001 compared with AAV-DIO-GFP vector (Student’s *t* test). Scale bars: 200 μm (A).

### Impact of USS length on Cre-independent expression of transgene

To evaluate the impact of the USS length on non-specific gene expression in the wild-type brain tissues, we generated AAV-DIO/3’USS-GFP vector with different sizes of the USS (750 bp for the controls and 553/285/103 bp for the shortening) (see Figure 4A for the structure and Figure S11 for location of restriction enzyme sites for size determination). These vectors (2.4 × 10^12^ genome copies/mL) were used for the intracranial injection into the striatum (1.0 μL/site × 4 sites) of wild-type Long Evans rats. Striatal sections were immunostained for GFP and typical images of the staining are shown in Figure 4B. The number of GFP^+^ cells showed a gradual increase along with the shortening of the USS (Figure 4C; one-way ANOVA, *F*_3,12_ = 218.396, *P* < 0.001). The cell number in AAV-DIO/3’USS(553)-GFP (2.25 ± 0.15 cells, n = 4 animals), AAV-DIO/3’USS(285)-GFP (3.06 ± 0.08 cells, n = 4 animals), and AAV-DIO/3’USS(103)-GFP (3.94 ± 0.15 cells, n = 4 animals) vectors was significantly larger than that in AAV-DIO/3’USS-GFP vector (not detected, n = 4 animals) (Bonferroni test, *P* < 0.001). These data highlight that the distance between the *lox*P and *lox*2272 sites at the unilateral side of inverted GFP cassette is important for the protection of non-specific transgene expression, and that 3’-USS with the size of 750 bp is at least necessary for intense protection.

**Figure 4.**
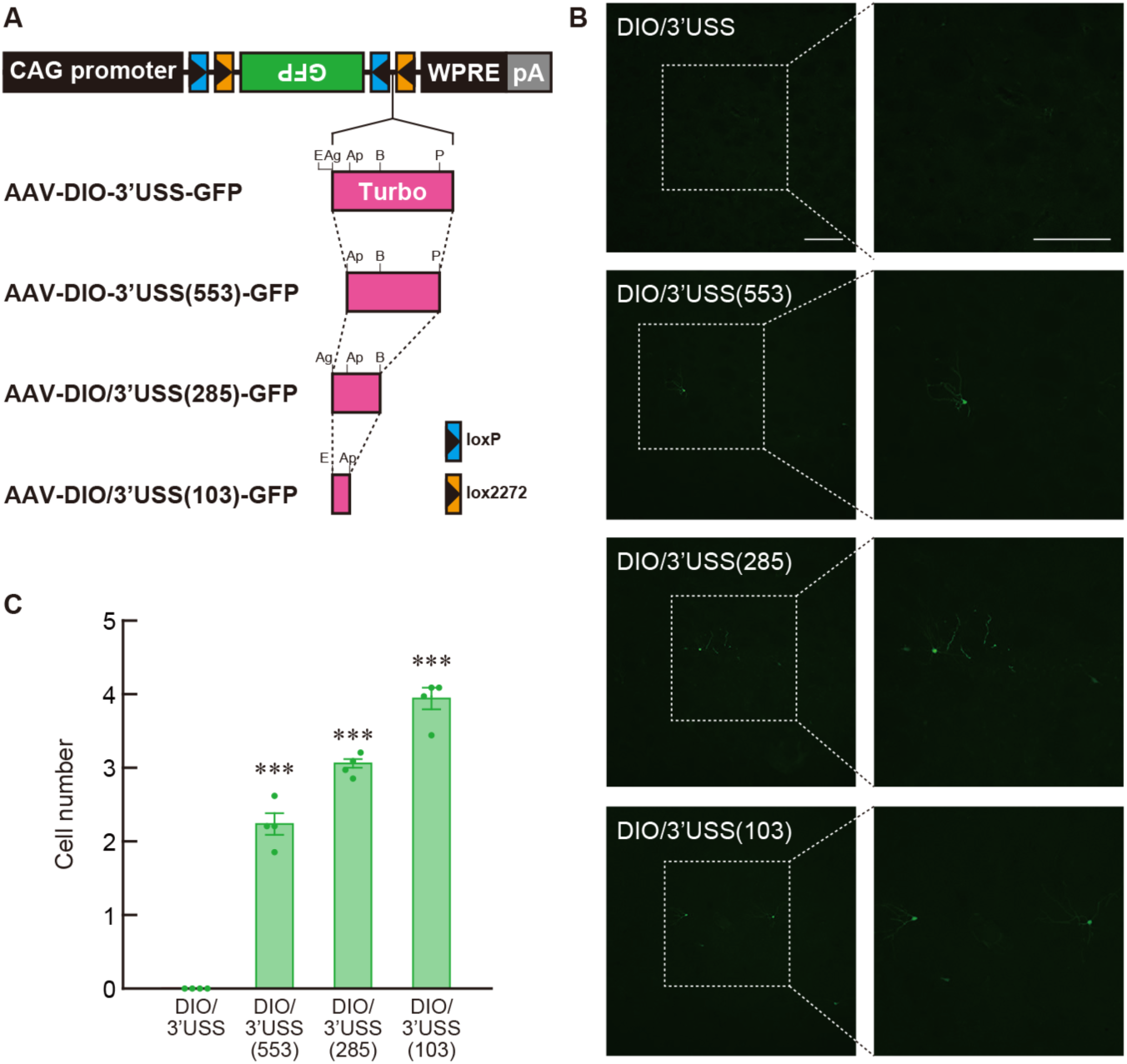
Influence of USS length on Cre-independent expression of transgene. (A) Genome structures of AAV-DIO/3’USS-GFP vectors with different sizes of the USS. The sequence encoding TurboFP635 (750 bp) in AAV-DIO/3’USS-GFP vector was replaced by its shorter fragments with 553, 285, or 103 bp. Restriction enzyme site abbreviations: Ag, *Age*I; Ap, *Apa*LI; B, *Bsu*36I; E, *Eco*47III; and P, *Psh*AI. pA: hGH polyadenylation signal. (B) GFP immunohistochemistry with striatal sections. Long Evans rats received the injection of four kinds of vectors (2.4 × 10^12^ copies/mL) into the striatum (0.1 μL/site × 4 sites), and striatal sections were immunostained for GFP. Typical immunostaining images are indicated for each vector. On the right side of the figure, magnified views of the squares in the left column of photos. (C) Shortening of 3’USS gradually increased the GFP^+^ cell number. Data are presented as mean ± SEM (n = 4 animals). Individual data are overlaid. \*\*\**P* < 0.001 compared with AAV-DIO/3’USS-GFP vector (Bonferroni test). Scale bars: 200 μm (B).

### Using the DIO/USS system for highly selective expression of other transgenes

To search whether the DIO/USS system is applicable for the selective expression of other transgenes, we converted the inverted transgene in the gene cassette of AAV-DIO-GFP or AAV-DIO/3’USS-GFP vector from GFP to a variant of chanelrhodopsin-2 fused to Venus (ChRWR/Venus) (Wang et al., 2009), a mutant form of human muscarinic M4 acetylcholine receptor connected to 2A-GFP (hM4Di/2A/GFP) (Kato et al., 2018) or a mutant form of red fluorescent protein (RFP; mCherry) (Shaner et al., 2004) (Figure 5A). The CAG promoter in these vectors was changed to the promoter of the gene encoding human elongation factor-1α (EF-1α) because of the shorter length of the EF-1α promoter (Kim et al., 1990), and eleven in-frame methionine codons in the TurboFP635 sequence were mutated to the termination codon TAG to prevent the synthesis of TurboFP635 protein or its fragments (see Figure S11). First, we checked non-specific expression of transgenes in the wild-type brain tissues. The AAV vectors with equivalent titers between the corresponding vectors (2.1 × 10^12^ genome copies/mL for AAV-DIO-ChRWR/Venus and AAV-DIO/3’USS- ChRWR/Venus, 2.4 × 10^12^ genome copies/mL for AAV-DIO-hMD4i/2A/GFP and AAV-DIO/3’USS- hMD4i/2A/GFP, and 2.2 × 10^12^ genome copies/mL for AAV-DIO- mCherry and AAV-DIO/3’USS-mCherry) were injected into the striatum (1.0 μL/site × 4 sites) of Long Evans rats. Striatal sections were immunostained with anti-GFP or anti-RFP antibody, and typical images of the staining are presented in Figure 5B. In the ChRWR/Venus transgene, the number of immune-positive cells in AAV-DIO/3’USS- ChRWR/Venus vector (0.47 ± 0.19 cells, n = 4 animals) showed a significant reduction as compared to that in AAV-DIO-ChRWR/Venus vector (6.28 ± 0.89 cells, n = 4 animals) (Figure 5C; Student’s *t* test, *t*_6_ = 6.390, *P* < 0.001). In the hM4Di/2A/GFP transgene, the immuno-positive cell number of in AAV-DIO/3’USS-hM4Di/2A/GFP vector (1.14 ± 0.26 cells, n = 4 animals) was significantly decreased compared to that in AAV-DIO-hM4Di/2A/GFP vector (5.13 ± 0.71 cells, n = 4 animals) (Figure 5C; Student’s *t* test, *t*_6_ = 5.236, *P* < 0.01). In the mCherry gene, the number of positive cells in AAV-DIO/3’USS-mCherry vector (1.05 ± 0.32 cells, n = 4 animals) was also significantly lower than that in AAV-DIO-mCherry vector (7.32 ± 0.51 cells, n = 4 animals) (Figure 5C; Student’s *t* test, *t*_6_ = 10.491, *P* < 0.001). These data indicate that the DIO system with 3’USS can suppress non-specific expression of other transgenes in wild-type brains.

**Figure 5.**
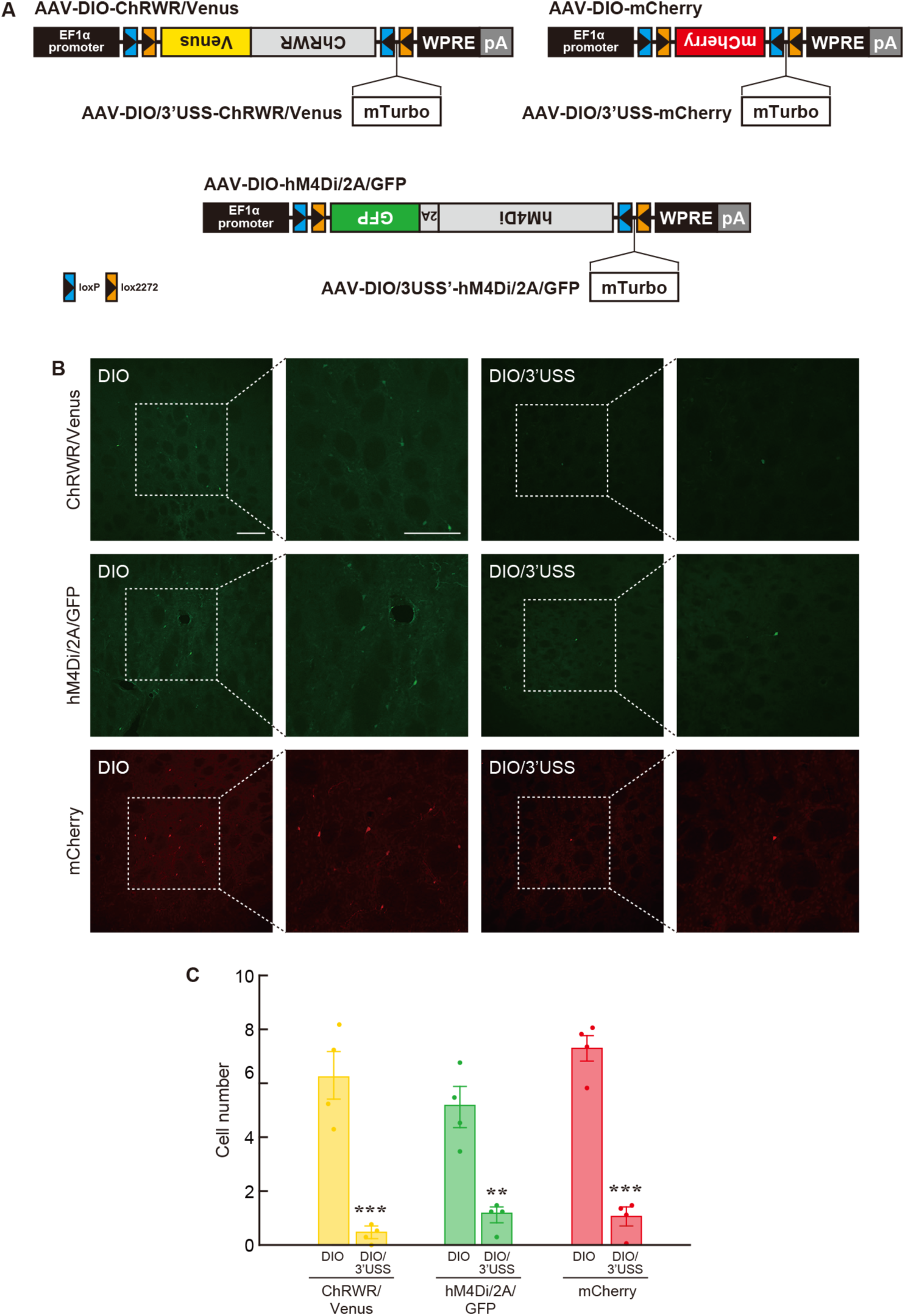
Silencing Cre-independent expression of other transgenes using the USS. (A) Genome structures of AAV-DIO/3’USS vectors for expression of ChRWR/Venus, hM4Di/2A/GFP, and mCherry transgenes. The mTurbo sequence, in which eleven in-frame methionine codons in the TurboFP635 sequence were substituted by termination codons, was used as the 3’USS. pA: hGH polyadenylation signal. (B) Immunostaining with anti-GFP or anti-RFP antibody with striatal sections. Long Evans rats received injections of AAV-DIO-ChRWR/Venus and AAV-DIO/3’USS-ChRWR/ Venus vectors (2.1 × 10^12^ copies/mL), AAV-DIO-hM4Di/2A/GFP and AAV-DIO/ 3’USS-hM4Di/2A/GFP vectors (2.4 × 10^12^ copies/mL), as well as AAV-DIO-mCherry and AAV-DIO/3’USS-mCherry vectors (2.2 × 10^12^ copies/mL) into the striatum (0.1 μL/site × 4 sites), and striatal sections were stained for GFP or RFP. Typical immunostaining images are presented for each vector. Right, Magnified views of the squares in the left photos. (C) Introduction of the 3’USS into the vectors suppresses the number of cells expressing the transgenes. Data are presented as mean ± SEM (n = 4 animals). Individual data are overlaid. \*\**P* < 0.01, \*\*\**P* < 0.001 compared with the corresponding AAV-DIO vectors (Student’s *t* test). Scale bars: 200 μm (B).

To validate the selectivity of Cre-dependent expression of other transgenes using the DIO/USS system, we studied the expression pattern of ChRWR/Venus transgene via injection with AAV-DIO-ChRWR/Venus and AAV-DIO/3’USS-ChRWR/Venus vectors (2.1 × 10^12^ genome copies/mL) into the striatum (1.0 μL/site × 4 sites) of PV-Cre knock-in rats. Striatal sections were stained by double immunohistochemistry with anti-PV and anti-GFP antibodies, and typical images of immunostaining are presented in Figure 6A. The percentage of the number of PV^+^ + ChRWR/Venus^+^ cells divided by that of PV^+^ cells was 78.45 ± 1.57% for AAV-DIO-ChRWR/Venus vector (n = 4 animals) and 79.22 ± 1.38% for AAV-DIO/3’USS-ChRWR/Venus vector (n = 4 animals) (Figure 6B; Student’s *t* test, *t*_6_ = 0.370, *P* = 0.724), indicating the efficient expression of ChRWR/Venus in PV-containing neurons in both vector-injected rats. In contrast, the percentage of the number of PV^+^ + ChRWR/Venus^+^ cells divided by that of ChRWR/Venus^+^ cells was significantly increased in AAV-DIO/3’USS-ChRWR/Venus vector (90.25 ± 0.33%) as compared to AAV-DIO-ChRWR/Venus vector (60.45 ± 0.89%), whereas the percentage of the number of PV^-^ + ChRWR/Venus^+^ cells divided by that of ChRWR/Venus^+^ cells was markedly decreased in AAV-DIO/3’USS- ChRWR/Venus vector (9.75 ± 0.33%) relative to AAV-DIO-ChRWR/Venus vector (39.55 ± 0.89%) (Figure 6C; Student’s *t* test, *t*_6_ = 31.257, *P* < 0.001). These results support enhanced selectivity of Cre-dependent expression of transgenes by the DIO system with 3’USS.

**Figure 6.**
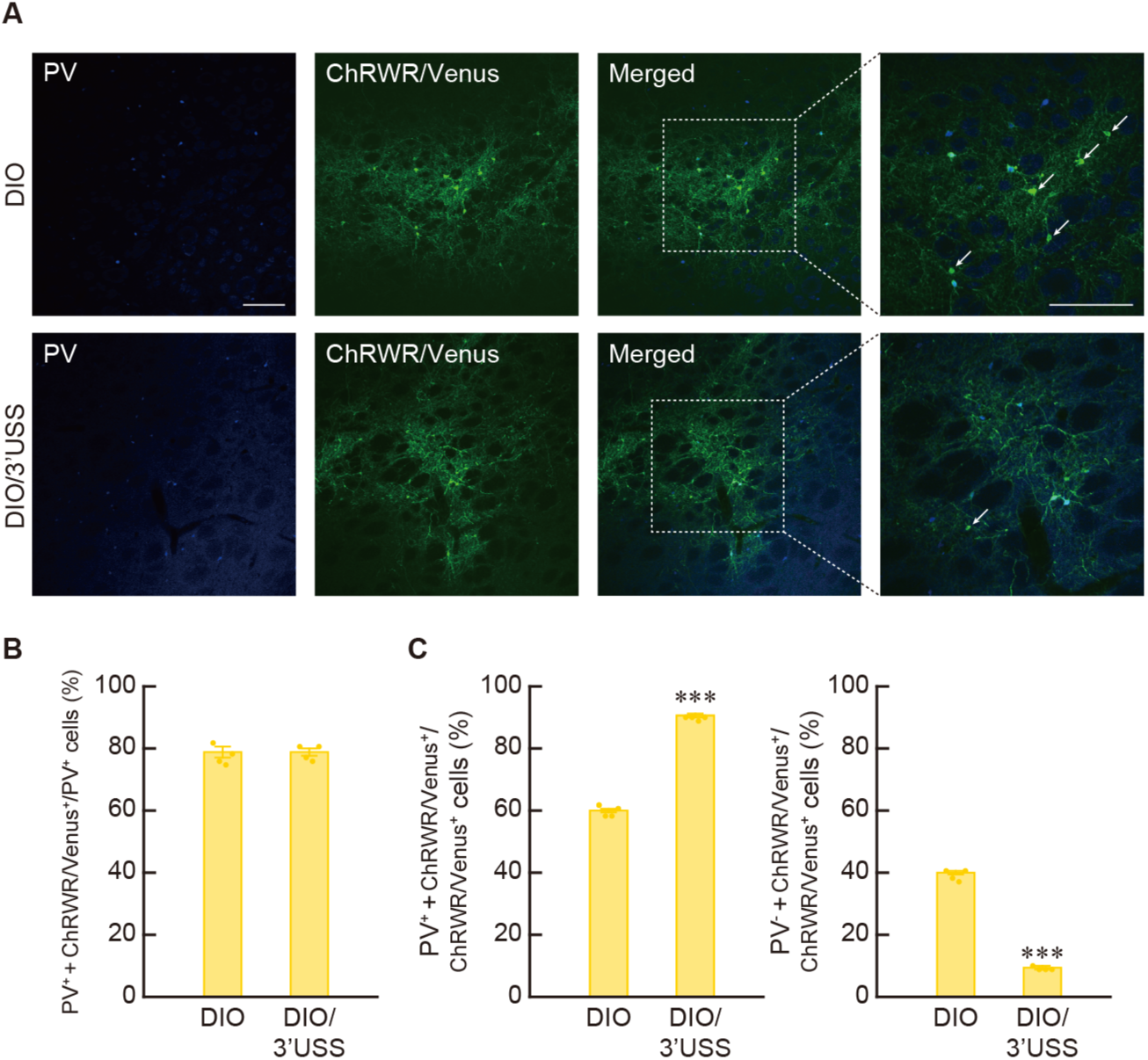
Enhanced selectivity of Cre-dependent expression of ChRWR/Venus through the DIO/USS system. (A) Double immunohistochemistry for PV and ChRWR/Venus with striatal sections. PV-Cre knock-in rats received the injection of AAV-DIO-ChRWR/Venus or AAV-DIO/3’USS- ChRWR/Venus vector (2.1 × 10^12^ copies/mL) into the striatum (0.1 μL/site × 4 sites), and sections through the striatum were used for double immunohistochemistry with anti-PV and anti-GFP antibodies. Typical images of double immunostaining are shown for each vector. Right, magnified views of the squares in the merged photos. Arrows indicate PV^-^/ChRWR/Venus^+^ cells. (B) High efficiency of Cre-dependent expression by the DIO/USS system. The percentage of the number of PV^+^/ChRWR/Venus^+^ cells to that of PV^+^ cells was calculated. (C) Enhanced selectivity of Cre-dependent expression by the DIO/USS system. The percentage of the number of PV^+^/ChRWR/Venus^+^ cells to that of ChRWR/Venus^+^ cells or the number of PV^-^/ChRWR/Venus ^+^ cells to that of ChRWR/Venus^+^ cells was calculated. Data are presented as mean ± SEM (n = 4 animals). Individual data are overlaid. \*\*\**P* < 0.001 compared with AAV-DIO-ChRWR/Venus vector (Student’s *t* test). Scale bars: 200 μm (A).

## DISCUSSION

A previous study reported that non-specific expression of transgenes in off-targets after treatment of AAV vectors carrying the DIO system are derived at least from infrequent transcription from inverted transgene and recombination events through short homologous sequences within recognition sites during the transfer plasmid amplification in bacterial cells (Fischer et al., 2019). In their study, bacterial recombination events were alleviated by decreasing sequence homology within the recognition sites, but the efficiency of Cre-mediated recombination was also deteriorated. To suppress transcription from the inverted transgene, the in-frame initiation codon was localized upstream of the recognition sites at the 5’-end of the transgene, in which Cre-mediated recombination deletes a gene cassette containing polyadenylation signals flanked by the two recognition sites in the same orientation. However, in this construct, an extra amino acid sequence was added in the N-terminal region of mature transgene products, which may interfere with processing or function of the transgene products. In the present study, although we did not detect recombination events of the transfer plasmid during its amplification in bacterial cells, we observed events in viral genome during production of AAV vector with the DIO system in cultured cells, which appeared to be generated through recombination between a pair of short nucleotide sequences within the *lox*P or *lox*2272 sites. The frequency of this recombination was markedly suppressed by introduction of the USS between double recognition sites at the unilateral side of the transgene, and the suppressive effect was greater at the 3’-end as compared to that at the 5’-end, which resulted in protection of non-specific transgene expression, improving the selectivity of Cre-dependent expression in target cell populations. Therefore, our DIO/USS system provides a useful strategy for highly selective Cre-dependent transgene expression with no influence on the efficiency of Cre-mediated recombination as well as the processing and function of transgene products.

Recombination events in AAV genome carrying the DIO sequence seemed to occur between a pair of short nucleotide sequences within the *lox*P or *lox*2272 sites during the vector production in cultured cells. When the plasmid constructs for AAV vector production are transfected into packaging cells, AAV DNA replication is initiated by cellular DNA polymerase activities together with Rep78 and Rep68 proteins provided from the RC gene plasmid, leading to the formation of double-stranded DNA replicative intermediates in monomer and dimer forms, from which single-stranded viral genomes with plus and minus polarities are synthesized and packaged into vector particles (Gonçalves, 2005). During viral DNA replication, recombination based on a short sequence homology may cause the reversion of the gene cassette in AAV genome with the DIO sequence. One possible mechanism that explains this reaction may involve homologous recombination, which mediates DNA repair for double-strand breaks (DSBs) (Li and Heyer, 2008). Although homologous recombination is generally considered to require long homologous sequences (Hockemeyer et al., 2009; Sommer et al., 2013), recent studies of gene editing with a single-strand template suggest that the recombination is also mediated via the homology of relatively shorter sequences (Paquet et al., 2016; Davis and Maizels, 2016). In addition, short homologous sequences of 5−25 bp are known to be implicated in microhomology-mediated end joining, which is another repair mechanism for DSBs (McVey and Lee, 2008; Truong et al., 2013). DSBs occur naturally during cell division when DNA replication forks encounter a blocking lesion, leading to fork stall and collapse (Shrivastav et al., 2008). Inverted repeats that potentially form secondary structures such as hairpin and cruciform structures appear to promote generation of DSBs at the site of repeats in eucaryotic cells (Lobachev et al., 2007). The inverted repeats at the *lox*P and *lox*2272 sites may produce secondary structures that interfere with progression of the replication forks, and then generate DSBs within these recognition sites, thereby leading to the reversion of transgene in AAV genome through short sequence homology-based recombination.

Introduction of the USS between the *lox*P and *lox*2272 sites at the 5’ or 3’-end of the inverted transgene in the DIO system effectively suppressed Cre-independent recombination in viral genome and non-specific expression of transgene in brain tissues. However, shortening of the USS between the two recognition sites increased the frequency of non-specific transgene expression, indicating the importance of the distance between the recognition sites for protection of non-specific gene expression. The size of 750 bp as 3’-USS was at least required for this protection. As aforementioned, secondary structures formed by inverted repeats in the two recognition sites may be involved in short sequence homology-based recombination in AAV genome carrying the DIO system. The USS introduction between the recognition sites is hypothesized to shift the secondary structures by distancing two pairs of inverted repeats at the *lox*P and *lox*2272 sites, and contribute to the suppressing effects against DSB formation and subsequent short sequence homology-based recombination. However, this hypothesis cannot explain the results showing that introduction of the USS into the 3’-end of the inverted gene cassette was more effective as compared to that into 5’-end of the cassette. Investigation of the molecular mechanism is needed that explains spacing effects on Cre-independent recombination events, including the stronger effects of 3’USS relative to 5’USS on the suppression of recombination.

Injection of AAV-DIO/3’USS-GFP vector into the brains of PV-Cre knock-in rats resulted in efficient expression of GFP in PV^+^ neurons and suppressed non-specific GFP expression in PV^-^ cells, resulting in the improved selectivity of Cre-dependent transgene expression. In contrast, the number of PV^-^/GFP^+^ cells in PV-Cre rats (2.53 ± 0.28 cells, n = 4 animals) was increased as compared to that of GFP^+^ cells obtained from the experiments, in which the same vector was injected into the wild-type brains (Figure 2E; 0.03 ± 0.03 cells, n = 4 animals), showing a significant difference between the two values (Student’s *t* test, *t*_6_ = 8.800, *P* < 0.001). We did not observe any Cre^+^ signals in PV^-^ neurons in the knock-in rat striatum in our double immunohistochemistry (see Figure S10). However, in these rats there may be a small number of cells expressing a low level of Cre protein that was not detectable in our experimental conditions, and these cells may express GFP thorough site-specific recombination.

In addition, a previous study reported that infrequent transcription from the inverted transgene in viral genome causes non-specific transgene expression in off-target cell populations (Fischer et al., 2019). We tested this possibility in our vector system by introducing a gene cassette containing rabbit *β*-globin gene polyadenylation signal in the reverse orientation downstream of hGH gene polyadenylation signal of the transfer plasmid to prevent transgene expression based on transcription in the reverse direction. After the vector injection into the rat striatum, the number of transgene-expressing cells was not different between the presence and absence of the polyadenylation signal in the reverse orientation in the vectors (Figure S12), excluding the possibility of gene expression based on transcription of the inverted transgene. These data support our observations that non-specific transgene expression after vector injection is mainly attributable to Cre-independent recombination in AAV genome with DIO sequence during vector production in packaging cells.

We successfully achieved the protection of Cre-independent recombination during AAV production and non-specific expression of transgene in off-targets, enhancing the selectivity of Cre-dependent gene expression by introducing the USS between the *lox*P and *lox*2272 sites at either the 5’- or 3’-end of the inverted transgene, in particular at the 3’-end with the more efficient protection. This DIO system with the USS was also applicable for the selective expression of different types of transgenes. Our DIO/USS system will provide a powerful tool for highly specific Cre-dependent transgene expression for the labeling of target cell populations and functional imaging, as well as the manipulation of the activity of these populations in the future.

## ACKNOWLEDGMENTS

This work was supported by: a grant-in-aid for Scientific Research (C) (MO20K06912) from the Japan Society for the Promotion of Science (S.K.); a grant-in-aid for Scientific Research on Transformative Research Areas (A) Adaptive Circuit Census (21H05244) from the Ministry of Education, Science, Sports, and Culture of Japan; and a grant-in-aid from the Japan Agency for Medical Research and Development under Grant Number (JP17dm0207052) (K.K.). The plasmid DNAs encoding ChRWR, hM4Di, and mCherry genes were kindly gifted from Drs. H. Yawo, B. Roth, and I. Wada, respectively. We are grateful to R. Fukabori, Y. Hashimoto, M. Kikuchi, and Y. Nakazato for their technical support during the animal experiments, and to T. Kobayashi for her helpful illustrations.

## AUTHOR CONTRIBUTIONS

N.M., S.K., and K.K. conceived the study, designed the experiments, and directed the project. N.M., S.K., K. T. and K.K. designed the vector expression strategy and generated the vector. N.M., Y.M., and T.M. produced and analyzed knock-in transgenic rats. N.M., S.K., and K. T. analyzed vector structure, and K.N. and M.S. performed intracranial injections and histological examinations. N.M., S.K., and K.K. wrote the paper. All authors discussed the results and implications, and commented on the manuscript at all stages.

## STAR★METHODS

### KEY RESOURCE TABLE

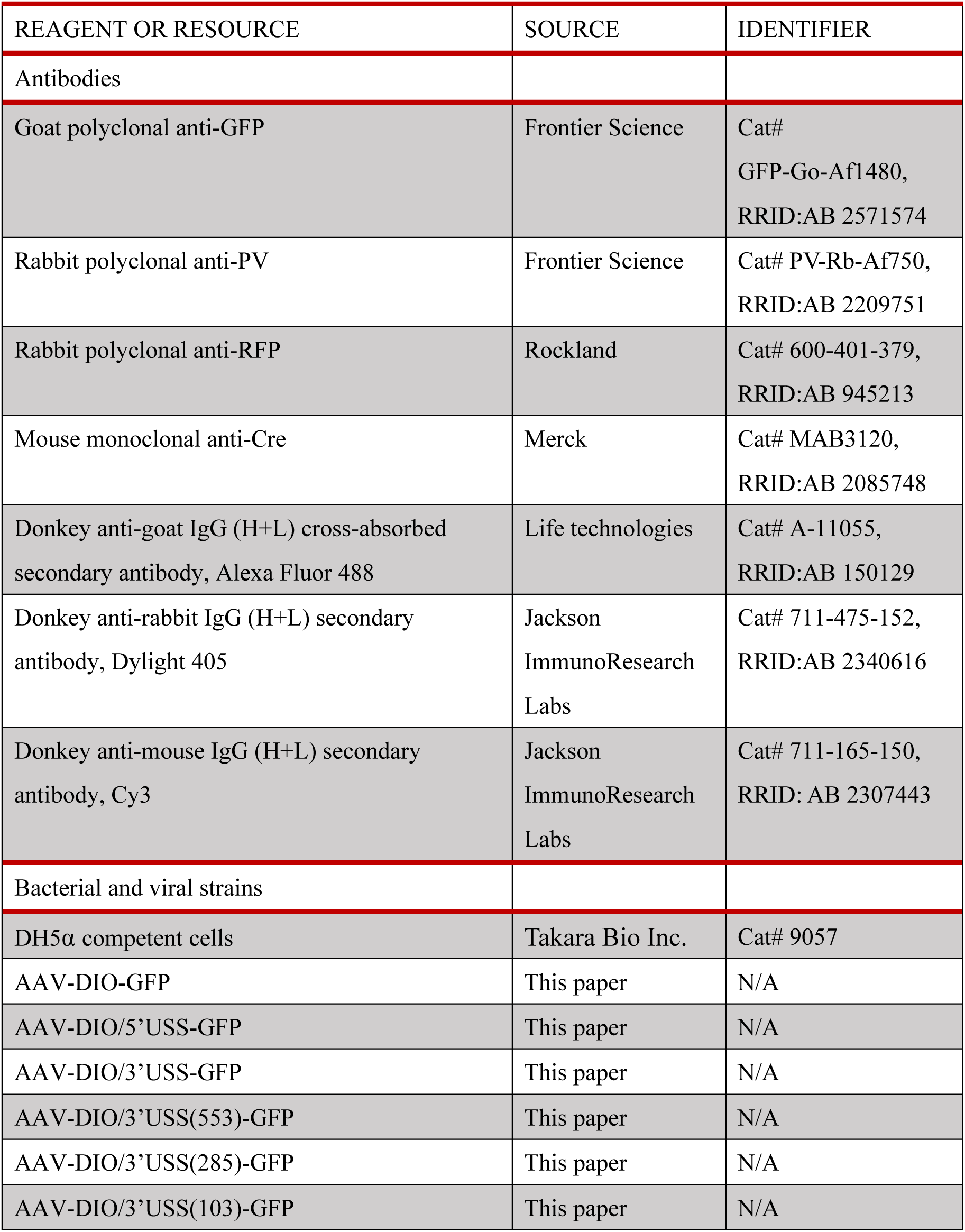
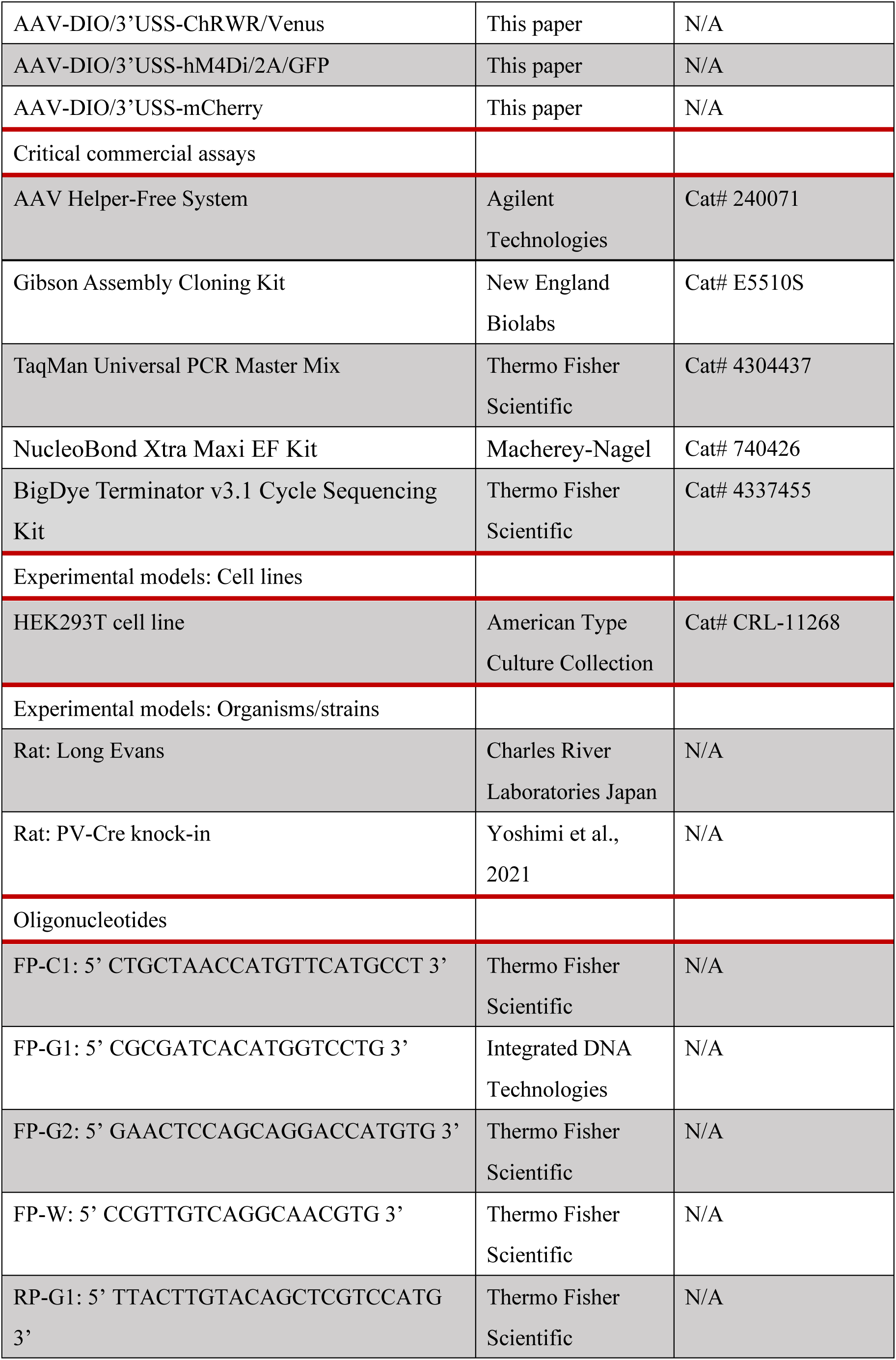
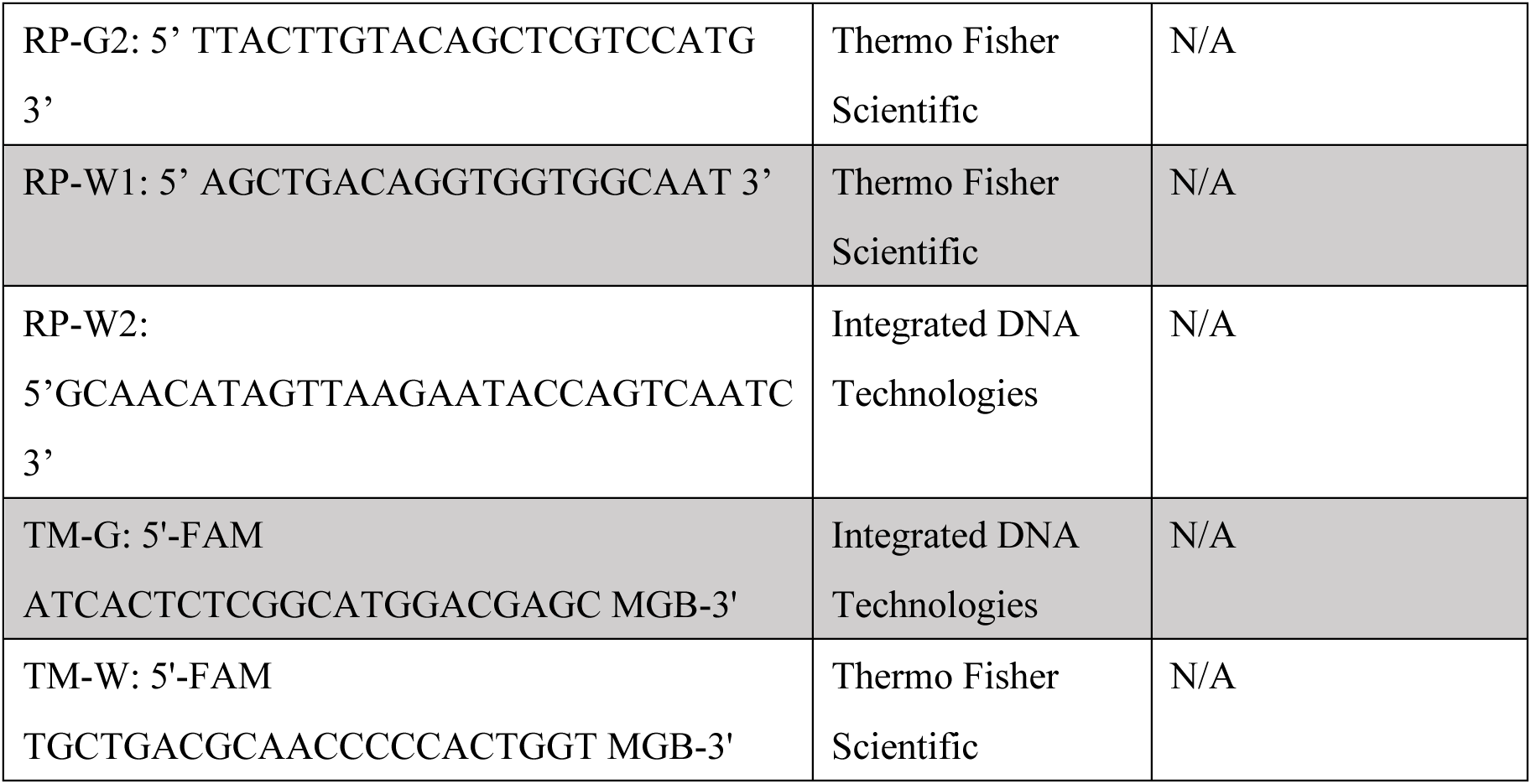

### RESOURCE AVAILABILITY

#### Lead contact

Further information and requests for resources and reagents should be directed to and will be fulfilled by the Lead Contact, Kazuto Kobayashi.

#### Material availability

Viral vectors and plasmids used in this study are available by direct distribution after completion of a standard MTA. PV-Cre knock-in rats are deposited to the National BioResource Project for the Rat in Japan (http://www.anim.med.kyoto-u.ac.jp/nbr/ Default. aspx).

#### Data and code availability

- Original data reported in this paper will be deposited at Mendeley Data after the acceptance of this study.
- This paper does not report original code.
- Any additional information required to reanalyze the data reported in this paper is available from the lead contact upon request.

### EXPERIMENTAL MODEL AND SUBJECT DETAILS

Animal care and handling procedures were conducted in accordance with the guidelines established by the Laboratory Animal Research Center of Fukushima Medical University. All procedures were approved by the Fukushima Medical University Institutional Animal Care and Use Committee. Rats were maintained at 22 ± 2°C and 60% humidity in a 12-hr light/12-hr dark cycle, and food and water were continuously available. Long Evans rats (Charles River Laboratories) and PV-Cre knock-in rats (Yoshimi et al., 2021) were used for the intracranial surgery. Both male and female rats at the age of 8–12 weeks were used for the experiments. HEK293T cells (American Type Culture Collection No. CRL-11268) were stored in frozen aliquots, and thawed and propagated in 10-cm dishes before use in viral production. Cells were cultured in Dulbecco’s modified Eagle’s medium (Sigma-Aldrich) containing 10% fetal bovine serum (Invitrogen) and supplemented with 2 mM glutaMax supplement (Gibco), and penicillin-streptomycin of 100 units/mL (Gibco) at 37°C with 5% CO_2_, and used for transfection.

### METHODS DETAILS

#### Plasmid construction

The transfer plasmid pAAV-DIO-GFP containing the gene cassette encoding GFP (Clontech) in the inverted orientation, which is flanked by two double recognition sites between the CAG promoter (Niwa et al., 1990) and WPRE sequence connected to hGH gene polyadenylation signal, was constructed using a Gibson Assembly Cloning Kit (New England Biolabs) (see Figure 1A). The gene cassette (750 bp) encoding TurboFP635 (Evrogen) was inserted between the *lox*P and *lox*2272 sites either at the 5’- or 3’-end of the inverted GFP sequence, resulting in the plasmids pAAV-DIO/5’USS-GFP and pAAV-DIO/3’USS-GFP (see Figure 2A). The TurboFP635 gene cassette in pAAV-DIO/3’USS-GFP was replaced by a 553-bp *Apa*LI/*Psh*AI, 285-bp *Eco*47III/*Bsu*36I, and 103-bp *Age*I/*Apa*LI DNA fragments of the cassette, resulting in pAAV-DIO/3’USS (553)-GFP, pAAV-DIO/3’USS (285)-GFP and pAAV-DIO/3’USS (103)-GFP, respectively (see Figure 4A). In the plasmid pAAV-DIO/3’USS-GFP, the CAG promoter was substituted by the promoter of the gene encoding human EF1α (Kim et al., 1990), eleven in-frame methionine codons in the TurboFP635 sequence were mutated to the termination codon TAG (termed mTurbo), and then the inverted GFP sequence was converted to the gene cassettes for ChRWR/Venus (Wang et al., 2009), hM4Di/2A/GFP (Kato et al., 2018), and mCherry (Shaner et al., 2004), designed the plasmids pAAV-EF1α-DIO/3’USS- ChRWR/Venus, pAAV-EF1α-DIO/3’USS-hM4Di/2A/GFP, and pAAV-EF1α-DIO/ 3’USS-mCherry, respectively (see Figure 5A).

#### Plasmid preparation

*E. coli* DH5α strain containing a mutation in the *Rec*A1 gene (Takara Bio Inc.) was used for DNA preparations. *E. coli* competent cells were transformed with appropriate amounts of plasmids. The colonies were picked up and cultured in a shaker at 160 rpm in 3 mL of LB containing 50 µg/mL ampicillin at 37°C for 8 h. Then, the medium was scaled up to 500 mL and continued to be cultured for 16 h. Plasmids were extracted and purified with a NucleoBond Xtra Maxi EF Kit (Macherey-Nagel).

#### Viral vector preparation

AAV vector serotype 2 was prepared based on AAV Helper-Free System (Agilent Technologies) as described (Kato et al., 2018). HEK293T cells (American Type Culture Collection) were transfected with the transfer gene plasmid, together with the RC gene and adeno-helper gene plasmids through the calcium phosphate precipitation method. The crude viral lysate was purified with two rounds of CsCl gradient ultracentrifugation at 100,000×*g* and 4°C for 23 h using 13.5-mL Quick-Seal tubes and a NVT65 rotor (Beckman Coulter). Each tube was placed gently and 500–750 µL of white turbid bands were collected. The solution was applied to three rounds of dialysis against 1 liter of PBS by using a Slide-A-Lyzer dialysis cassette (MWCO10,000; Thermo Fisher Scientific). The dialyzed vector solution was finally concentrated by centrifugation through a Vivaspin Turbo filter (Sartorius). The viral genome titer was determined by quantitative PCR using the TaqMan System (Thermo Fisher Scientific) with the following primer sets and probes. Total vector titer was measured with the FP-W/RP-W1 primer set, which amplifies a part of WPRE, and the probe TM-W. Recombinant vector titer was measured with the FP-G1/RPW2 primer set, which amplifies the sequence containing the 3’ part of GFP transgene, and the probe TM-G.

#### PCR amplification

PCR assays were carried out in the reaction mixture (10 μL) containing 1.5 units rTaq DNA polymerase, 10 pmol each of forward and reverse primer pair, 2.5 nmol each dNTP mixture in PCR buffer (10 nmol Tris-HCl; pH 8.3, 50 nmol KCl, and 1.5 nmol MgCl_2_) (Takara Bio Inc.). The amplifications were performed with 25 cycles of denaturation at 96°C for 10 s, annealing at 58°C for 20 s, and extension at 72°C for 75 s. The PCR products were analyzed by electrophoresis in 1% agarose gels containing ethidium bromide with 100-bp ladder marker (Takara Bio Inc.). The DNA fragments were excised from the gel and eluted into 10 mM Tris-HCl buffer (pH8.0) containing 1 mM EDTA by using QIAGEN II Gel Extraction Kit (QIAGEN).

#### Nucleotide sequence analysis

Dye-terminator sequencing was carried out in the reaction mixture (10 μL) containing a 3 pmol specific primer by using a BigDye Terminator v3.1 Cycle Sequencing Kit (Thermo Fisher Scientific). The chain terminations were performed with 25 cycles of denaturation at 96°C for 10 s, annealing at 50°C for 5 s, and extension and termination at 60°C for 3 min. The extension products were collected by high-speed centrifugation after ethanol addition, dissolved in the Hi-Di formamide (Thermo Fisher Scientific), and then analyzed by the capillary electrophoresis instrument (3500 Genetic Analyzer with a data collection software; Thermo Fisher Scientific). The nucleotide sequences were determined by using sequencing analysis software (Thermo Fisher Scientific).

#### Intracranial surgery

The surgery was conducted under isoflurane (4% induction and 1.5% maintenance) anesthesia, and AAV vectors were introduced into the dorsal striatum (1.0 μL/site, 4 sites) through a glass microinjection capillary connected to a microinfusion pump (ESP-32; Eicom). The anteroposterior, mediolateral, and dorsoventral coordinates (mm) from bregma and dura were 1.5/3.0/3.0 (site 1), 1.5/3.0/3.5 (site 2), 0.5/3.0/3.0 (site 3), and 0.5/3.0/3.5 (site 4) according to an atlas of the rat brain (Paxinos and Watson, 2005). Injection was performed at a constant flow rate of 0.1 μL/min.

#### Immunohistochemistry

Rats were anesthetized with 4% isoflurane and perfused transcardially with PBS, followed by fixation with 4% paraformaldehyde in 0.1 M phosphate buffer (pH 7.4). Fixed brains were cut into sections (30-μm thick) through a coronal plane with a cryostat. For immunofluorescence histochemistry, sections were incubated with anti-GFP antibody (goat, 1 μg/mL; Frontier Science), anti-PV antibody (rabbit, 1 μg/mL; Frontier Science), and anti-RFP (rabbit, 1 μg/mL; Rockland), and then with species-specific secondary antibodies conjugated to DyLight 405 (Jackson ImmunoResearch Labs), Alexa 488 (Life technologies), or Cy3 (Jackson ImmunoResearch Labs). Fluorescent images were obtained with a confocal laser-scanning microscope (Nikon A1) equipped with proper filter cube specifications.

#### Cell counts

Eight sections through the dorsal striatum along with the anteroposterior coordinates (mm) between 1.68 and 0.48 from bregma were prepared from each rat, and the number of immuno-positive cells in ROIs (1.0 × 1.0 mm) in the striatum was counted (Figure S6). The average of stained cell number per section was calculated.

### QUANTIFICATION AND STASTICAL ANALYSIS

#### Statistical analysis

All data are presented as mean ± SEM. For multiple comparisons, one-way ANOVA was applied followed by *post hoc* Bonferroni test. An unpaired Student’s *t* test was also used. All statistical details of the experiments can be found in the Results section. Differences were considered statistically significant at *P* < 0.05.

### SUPPLEMENTAL INFORMATION

Supplemental information is attached as another file.

## REFERENCES

Abremski, K., Hoess, R., and Stenberg, N. (1983). Studies on the properties of P1 site-specific recombination: evidence for topologically unlinked products following recombination. Cell 32, 1301–1311. https://doi.org/10.1016/0092-8674(83)90311-2.

Aelvoet, S.A., Pascual-Brazo, J., Libbrecht, S., Reumers, V., Gijsbers, R., Van den Haute, C., and Baekelandt, V. (2015). Long-term fate mapping using conditional lentiviral vectors reveals a continuous contribution of radial glia-like cells to adult hippocampal neurogenesis in mice. PLoS One 10, e0143772. https://doi.org/10.1371/journal.pone.0143772.

Arguello, A.A., Richardson, B.D., Hall, J.L., Wang, R., Hodges, M.A., Mitchell, M.P., Stuber, G.D., Rossi, D.J., and Fuchs, R.A. (2017). Role of a lateral orbital frontal cortex-basolateral amygdala circuit in cue-induced cocaine-seeking behavior. Neuropsychopharmacology 42, 727–735. https://doi.org/10.1038/npp.2016.157.

Davis, L., and Maizels, N. (2016). Two distinct pathways support gene correction by single-stranded donors at DNA nicks. Cell Rep. 17, 1872–1881. https://doi.org/10.1016/j.celrep.2016.10.049.

Fischer, K.B., Collins, H.K., and Callaway, E.M. (2019). Sources of off-target expression from recombinase-dependent AAV vectors and mitigation with cross-over insensitive ATG-out vectors. Proc. Natl. Acad. Sci. USA 116, 27001–27010. https://doi.org/10.1073/pnas.1915974116.

Gonçalves, M.A. (2005). Adeno-associated virus: from defective virus to effective vector. Virol. J. 2, 43. https://doi.org/10.1186/1743-422X-2-43.

Gu, H., Zou, Y.R., and Rajewsky, K. (1993). Independent control of immunoglobulin switch recombination at individual switch regions evidenced through Cre-*loxP*-mediated gene targeting. Cell 73, 1155–1164. https://doi.org/10.1016/0092-8674(93)90644-6.

Hockemeyer, D., Soldner, F., Beard, C., Gao, Q., Mitalipova, M., DeKelver, R.C., Katibah, G.E., Amora, R., Boydston, E.A., Zeitler, B., et al. (2009). Efficient targeting of expressed and silent genes in human ESCs and iPSCs using zinc finger nucleases. Nat. Biotechnol. 27, 851–857. https://doi.org/10.1038/nbt.1562.

Hoess, R.H., Ziese, M., and Sternberg, N. (1982). P1 site-specific recombination: Nucleotide sequence of the recombining sites. Proc. Natl. Acad. Sci. USA 79, 3398–3402. https://doi.org/10.1073/pnas.79.11.3398.

Kato, S., Fukabori, R., Nishizawa, K., Okada, K., Yoshioka, N., Sugawara, M., Maejima, Y., Shimomura, K., Okamoto, M., Eifuku, S., et al. (2018) Action selection and flexible switching controlled by the intralaminar thalamic neurons. Cell Rep. 22, 2370–2382. https://doi.org/10.1016/j.cellrep.2018.02.016.

Kim, D.W., Uetsuki, T., Kaziro, Y., Yamaguchi, N., and Sugano, S. (1990). Use of the human elongation factor 1 alpha promoter as a versatile and efficient expression system. Gene 91, 217–223. https://doi.org/10.1016/0378-1119(90)90091-5.

Kozorovitskiy, Y., Saunders, A., Johnson, C.A., Lowell, B.B., and Sabatini, B.L. (2012). Recurrent network activity drives striatal synaptogenesis. Nature 485, 646–650. https://doi.org/10.1038/nature11052.

Lee, G., and Saito, I. (1998) Role of nucleotide sequences of loxP spacer region in Cre-mediated recombination. Gene 216, 55–65. https://doi.org/10.1016/s0378-1119(98)00325-4.

Li, X., and Heyer, W.D. (2008). Homologous recombination in DNA repair and DNA damage tolerance. Cell Res. 18, 99–113. https://doi.org/10.1038/cr.2008.1.

Lobachev, K.S., Rattray, A., and Narayanan, V. (2007). Hairpin- and cruciform-mediated chromosome breakage: causes and consequences in eukaryotic cells. Front. Biosci. 12, 4208–4220. https://doi.org/10.2741/2381.

McVey, M. and Lee, S.E. (2008). MMEJ repair of double-strand breaks (director’s cut): deleted sequences and alternative endings. Trends Genet. 24, 529–538. https://doi.org/10.1016/j.tig.2008.08.007.

Miyamichi, K., Shlomai-Fuchs, Y., Shu, M., Weissbourd, B.C., Luo, L., and Mizrahi, A. (2013). Dissecting local circuits: Parvalbumin interneurons underlie broad feedback control of olfactory bulb output. Neuron 80, 1232–1245. https://doi.org/10.1016/j.neuron.2013.08.027.

Niwa, H., Yamamura, K., and Miyazaki, J. (1991). Efficient selection for high-expression transfectants with a novel eukaryotic vector. Gene. 108, 193–199. https://doi.org/10.1016/0378-1119(91)90434-d.

Nonomura, S., Nishizawa, K., Sakai, Y., Kawaguchi, Y., Kato, S., Uchigashima, M., Watanabe, M., Yamanaka, K., Enomoto, K., Chiken, S. et al. (2018). Monitoring and updating of action selection for goal-directed behavior through the striatal direct and indirect pathways. Neuron 99, 1302–1314.e5. https://doi.org/10.1016/j.neuron.2018.08.002.

O’Connor, D.H., Hires, S.A., Guo, Z.V., Li, N., Yu, J., Sun, Q.Q., Huber, D., and Svoboda, K. (2013). Neural coding during active somatosensation revealed using illusory touch. Nat. Neurosci. 16, 958–965. https://doi.org/10.1038/nn.3419.

Paquet, D., Kwart, D., Chen, A., Sproul, A., Jacob, S., Teo, S., Olsen, K.M., Gregg, A., Noggle, S., and Tessier-Lavigne, M. (2016). Efficient introduction of specific homozygous and heterozygous mutations using CRISPR/Cas9. Nature 533, 125–129. https://doi.org/10.1038/nature17664.

Paxinos, G., and Watson, C. The Rat Brain in Stereotaxic Coordinates, Ed 6. Sydney: Academic Press (2007).

Sauer, B. (1987). Functional expression of the cre-lox site-specific recombination system in the yeast Saccharomyces cerevisiae. Mol. Cell Biol. 7, 2087–2096. https://doi.org/10.1128/mcb.7.6.2087-2096.1987.

Sauer, B., and Henderson, N. (1988). Site-specific DNA recombination in mammalian cells by the Cre recombinase of bacteriophage P1. Proc. Natl. Acad. Sci. USA 85, 5166–5170. https://doi.org/10.1073/pnas.85.14.5166.

Saunders, A., Johnson, C.A., and Sabatini, B.L. (2012). Novel recombinant adeno-associated viruses for Cre activated and inactivated transgene expression in neurons. Front. Neural Circuits 6, 47. https://doi.org/10.3389/fncir.2012.00047.

Schnütgen, F., Doerflinger, N., Calléja, C., Wendling, O., Chambon, P., and Ghyselinck, N.B. (2003). A directional strategy for monitoring Cre-mediated recombination at the cellular level in the mouse. Nat. Biotechnol. 21, 562–565. https://doi.org/10.1038/nbt811.

Shaner, N.C., Campbell, R.E., Steinbach, P.A., Giepmans, B.N., Palmer, A.E., and Tsien, R.Y. (2004). Improved monomeric red, orange and yellow fluorescent proteins derived from Discosoma sp. red fluorescent protein. Nat. Biotechnol. 22, 1567–1572. https://doi.org/10.1038/nbt1037.

Shcherbo, D., Merzlyak, E.M., Chepurnykh, T.V., Fradkov, A.F., Ermakova, G.V., Solovieva, E.A., Lukyanov, K.A., Bogdanova, E.A., Zaraisky, A.G., Lukyanov, S., et al. (2007). Bright far-red fluorescent protein for whole-body imaging. Nat Methods. 4, 741–746. https://doi.org/10.1038/nmeth1083.

Shrivastav, M., De Haro, L.P., and Nickoloff, J.A. (2008). Regulation of DNA double-strand break repair pathway choice. Cell Res. 18, 134–147. https://doi.org/10.1038/cr.2007.111.

Sohal, V.S., Zhang, F., Yizhar, O., and Deisseroth, K. (2009). Parvalbumin neurons and gamma rhythms enhance cortical circuit performance. Nature 459, 698–702. https://doi.org/10.1038/nature07991.

Sommer, D., Peters, A., Wirtz, T., Mai, M., Ackermann, J., Thabet, Y., Schmidt, J., Weighardt, H., Wunderlich, F.T., Degen, J., et al. (2014). Efficient genome engineering by targeted homologous recombination in mouse embryos using transcription activator-like effector nucleases. Nat. Commun. 5, 3045. https://doi.org/10.1038/ncomms4045.

Srinivasan, R., Lu, T.Y., Chai, H., Xu, J., Huang, B.S., Golshani, P., Coppola, G., and Khakh, B.S. (2016) New transgenic mouse lines for selectively targeting astrocytes and studying calcium signals in astrocyte processes in situ and in vivo. Neuron 92, 1181–1195. https://doi.org/10.1016/j.neuron.2016.11.030.

Sternberg, N., and Hamilton, D. (1981). Bacteriophage P1 site-specific recombination. I. Recombination between loxP sites. J. Mol. Biol. 150, 467–486. https://doi.org/10.1016/0022-2836(81)90375-2.

Truong, L.N., Li, Y., Shi, L.Z., Hwang, P.Y.H., He, J., Wang, H., Razavian, N., Berns, M.W., and Wu, X. (2013). Microhomology-mediated end joining and homologous recombination share the initial end resection step to repair DNA double-strand breaks in mammalian cells. Proc. Natl. Acad. Sci. USA 110, 7720–7725. https://doi.org/10.1073/pnas.1213431110.

Tschida, K., Michael, V., Takatoh, J., Han, B.X., Zhao, S., Sakurai, K., Mooney, R., and Wang, F. (2019). A specialized neural circuit gates social vocalizations in the mouse. Neuron 103, 459–472.e4. https://doi.org/10.1016/j.neuron.2019.05.025.

Voziyanov, Y., Pathania, S., and Jayaram, M. (1999). A general model for site-specific recombination by the integrase family recombinases. Nucleic Acids Res. 27, 930–941. https://doi.org/10.1093/nar/27.4.930.

Wang, H., Sugiyama, Y., Hikima, T., Sugano, E., Tomita, H., Takahashi, T., Ishizuka, T., and Yawo, H. (2009). Molecular determinants differentiating photocurrent properties of two channelrhodopsins from chlamydomonas. J. Biol. Chem. 284, 5685–5696. https://doi.org/10.1074/jbc.M807632200.

Woods, N.I., Stefanini, F., Apodaca-Montano, D.L., Tan, I.M.C., Biane, J.S., and Kheirbek, M.A. (2020). The dentate gyrus classifies cortical representations of learned stimuli. Neuron 107, 173–184.e6. https://doi.org/10.1016/j.neuron.2020.04.002.

Yoshimi, K., Oka, Y., Miyasaka, Y., Kotani, Y., Yasumura, M., Uno, Y., Hattori, K., Tanigawa, A., Sato, M., Oya, M., et al. (2021). Combi-CRISPR: combination of NHEJ and HDR provides efficient and precise plasmid-based knock-ins in mice and rats. Hum. Genet. 140, 277–287. https://doi.org/10.1007/s00439-020-02198-4.

