## Supplemental Information for "Highly selective transgene expression through double-floxed inverted orientation system by using a unilateral spacer sequence"

### SUPPLEMENTAL FIGURES

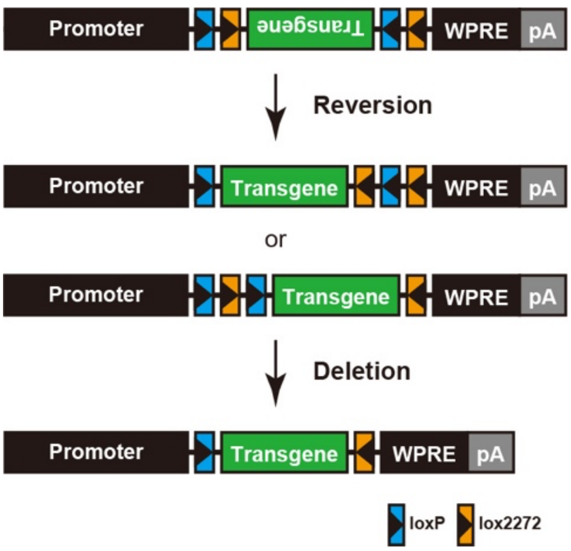


**Figure S1. Two-step Cre-dependent recombination through the DIO system**
A gene cassette in the inversed orientation is reversed through the first Cre-mediated recombination between two *lox*P sites or two *lox*2272 sites, which are located at both ends of the cassette in the opposite direction with an alternate order. Then, the second Cre-mediated recombination removes an intervening sequence flanked by two *lox*P sites or two *lox*2272 sites in the same direction, which are positioned either at 5' or 3'-end of the cassette. pA: polyadenylation signal.

pAAV-DIO-GFP

```
                 → CAG promoter
1     GCGGCCGCAC GCGTCGACAT TGATTATTGA CTAGTTATTA ATAGTAATCA ATTACGGGGT CATTAGTTCA TAGCCCATAT ATGGAGTTCC GCGTTACATA   100
        NotI       PstI
101   ACTTACGGTA AATGGCCCGC CTGGCTGACC GCCCAACGAC CCCCGCCCAT TGACGTCAAT AATGACGTAT GTTCCCATAG TAACGCCAAT AGGGACTTTC   200
201   CATTGACGTC AATGGGTGGA GTATTTACGG TAAACTGCCC ACTTGGCAGT ACATCAAGTG TATCATATGC CAAGTACGCC CCCTATTGAC GTCAATGACG   300
301   GTAAATGGCC CGCCTGGCAT TATGCCCAGT ACATGACCTT ATGGGACTTT CCTACTTGGC AGTACATCTA CGTATTAGTC ATCGCTATTA CCATGGTCGA   400
401   GGTGAGCCCC ACGTTCTGCT TCACTCTCCC CATCTCCCCC CCCTCCCCAC CCCCAATTTT GTATTTATTT ATTTTTTAAT TATTTTGTGC AGCGATGGGG   500
501   GCGGGGGGGG GGGGGGGGCG CGCGCCAGGC GGGGCGGGGC GGGGCGAGGG GCGGGGCGGG GCGAGGCGGA GAGGTGCGGC GGCAGCCAAT CAGAGCGGCG   600
                                                                                     * → transcription start
601   CGCTCCGAAA GTTTCCTTTT ATGGCGAGGC GGCGGCGGCG GCGGCCCTAT AAAAAGCGAA GCGCGCGGCG GGCGGGAGTC GCTGCGCGCT GCCTTCGCCC   700
                                                                 ▼
701   CGTGCCCCGC TCCGCCGCCG CCTCGCGCCG CCCGCCCCGG CTCTGACTGA CCGCGTTACT CCCACAGgtg agcgggcggg acggcccttc tcctccgggc   800
801   tgtaattagc gcttggttta atgacggctt gtttcttttc tgtggctgcg tgaaagcctt gaggggctcc gggagggccc tttgtgcggg gggagcggct   900
901   cggggggtgc gtgcgtgtgt gtgtgcgtgg ggagcgccgc gtgcggctcc gcgctgcccg gcggctgtga gcgctgcggg cgcggcgcgg ggctttgtgc   1000
1001  gctccgcagt gtgcgcgagg ggagcgcggc cgggggcggt gccccgcggt gcgggggggg ctgcgagggg aacaaaggct gcgtgcgggg tgtgtgcgtg   1100
1101  gggggtgag caggggtgt gggcgcgtcg gtcgggctgc aacccccccct gcacccccct ccccgagttg ctgagcacgg cccggcttcg ggtgcggggc   1200
1201  tccgtacggg gcgtggcgcg gggctcgccg tgccgggcgg ggggtggcgg caggtggggg tgccgggcgg ggcggggccg cctcgggccg gggagggctc   1300
1301  gggggagggg cgcggcggcc cccggagcgc cggcggctgt cgaggcgcgg cgagccgcag ccattgcctt ttatggtaat cgtgcgagag ggcgcaggga   1400
1401  cttcctttgt cccaaatctg tgcggagccg aaatctggga ggcgccgccg caccccctct agcgggcgcg gggcgaagcg gtgcggcgcc ggcaggaagg   1500
1501  aaatgggcgg ggagggcctt cgtgcgtcgc cgcgccgccg tccccttctc cctctccagc ctcggggctg tccgcggggg gacggctgcc ttcggggggg   1600
                                                                                                    ▼
1601  acggggcagg gcggggttcg gcttctggcg tgtgaccggc ggctctagag cctctgctaa ccatgttcat gccttcttct ttttcctaca gCTCCTGGGC   1700
1701  AACGTGCTGG TTATTGTGCT GTCTCATCAT TTTGGCAAAG AATTCATAAC TTCGTATAGC ATACATTATA CGAAGTTATG CAGAATGGTA GCTGGATTGT   1800
                                                 EcoRI               loxP        term
1801  AGCTGCTATT AGCAATATGA AACCTCTTAA TAACTTCGTA TAGGATACTT TATACGAAGT TATGGATCCT TACTTGTACA GCTCGTCCAT GCCGAGAGTG   1900
                                                   lox2272           BamHI
1901  ATCCCGGCGG CGGTCACGAA CTCCAGCAGG ACCATGTGAT CGCGCTTCTC GTTGGGGTCT TTGCTCAGGG CGGACTGGGT GCTCAGGTAG TGGTTGTCGG   2000
2001  GCAGCAGCAC GGGGCCGTCG CCGATGGGGG TGTTCTGCTG GTAGTGGTCG GCGAGCTGCA CGCTGCCGTC CTCGATGTTG TGGCGGATCT TGAAGTTCAC   2100
2101  CTTGATGCCG TTCTTCTGCT TGTCGGCCAT GATATAGACG TTGTGGCTGT TGTAGTTGTA CTCCAGCTTG TGCCCCAGGA TGTTGCCGTC CTCCTTGAAG   2200
2201  TCGATGCCCT TCAGCTCGAT GCGGTTCACC AGGGTGTCGC CCTCGAACTT CACCTCGGCG CGGGTCTTGT AGTTGCCGTC GTCCTTGAAG AAGATGGTGC   2300
2301  GCTCCTGGAC GTAGCCTTCG GGCATGGCGG ACTTGAAGAA GTCGTGCTGC TTCATGTGGT CGGGGTAGCG GCTGAAGCAC TGCACGCCGT AGGTCAGGGT   2400
2401  GGTCACGAGG GTGGGCCAGG GCACGGGCAG CTTGCCGGTG GTGCAGATGA ACTTCAGGGT CAGCTTGCCG TAGGTGGCAT CGCCCTCGCC CTCGCCGGAC   2500
2501  ACGCTGAACT TGTGGCCGTT TACGTCGCCG TCCAGCTCGA CCAGGATGGG CACCACCCCG GTGAACAGCT CCTCGCCCTT GCTCACCATG GTGGCAAGCT   2600
                                                                                           GFP ←
2601  TGCCTCGAGC AGCGCTGCTC GAGAGATCTA TAACTTCGTA TAATGTATGC TATACGAAGT TATTTGCCTT AACCCAGAAA TTATCACTGT TATTCTTTAG   2700
                             BglII              loxP                → WPRE
2701  AATGGTGCAA AGAATAACTT CGTATAAAGT ATCCTATACG AAGTTATGCG GCCGCATCGA Ttaatcaacc tctggattac aaaatttgtg aaagattgac   2800
                        lox2272                     NcoI     ClaI
2801  tggtattctt aactatgttg ctccttttac gctatgtgga tacgctgctt taatgccttt gtatcatgct attgcttccc gtatggcttt cattttctcc   2900
2901  tccttgtata aatcctggtt gctgtctctt tatgaggagt tgtggcccgt tgtcaggcaa cgtggcgtgg tgtgcactgt gtttgctgac gcaaccccca   3000
3001  ctggttgggg cattgccacc acctgtcagc tcctttccgg gactttcgct ttccccctcc ctattgccac ggcggaactc atcgccgcct gccttgcccg   3100
3101  ctgctggaca ggggctcggc tgttgggcac tgacaattcc gtggtgttgt cggggaagct gacgtccttt ccatggctgc tcgcctgtgt tgccacctgg   3200
3201  attctgcgcg ggacgtcctt ctgctacgtc ccttcggccc tcaatccagc ggaccttcct tcccgcggcc tgctgccggc tctgcggcct cttccgcgtc   3300
                                                                   → hGH gene
3301  ttcgccttcg ccctcagacg agtcggatct ccctttgggc cgcctccccg catctgatct acgggtggca tccctgtgac ccctccccag tgcctctcct   3400
3401  ggccctggaa gttgccactc cagtgcccac cagccttgtc ctaataaaat taagttgcat cattttgtct gactaggtgt ccttctataa tattatgggg   3500
                                                pA
3501  tggagggggg tggtatggag caaggggcaa gttgggaaga caacctgtag ggcctgcggg gtctattggg aaccaagctg gagtgcagtg gcacaatctt   3600
3601  ggctcactgc aatctccgcc tcctgggttc aagcgattct cctgcctcag cctcccgagt tgttgggatt ccaggcatgc atgaccaggc tcagctaatt   3700
3701  tttgtttttt tggtagagac ggggtttcac catattggcc aggctggtct ccaactccta atctcaggtg atctacccac cttggcctcc caaattgctg   3800
3801  ggattacagg cgtgaaccac tgctcccttc cctgtcctt
```

**Figure S2. Nucleotide sequence of pAAV-DIO-GFP transfer plasmid**

Asterisk indicates the transcription initiation site, and arrowheads show the splice donor and acceptor sites. The inverted GFP coding sequence, *lox*P site, and *lox*2272 site are shown in green, blue, and orange, respectively. pA: hGH polyadenylation signal.

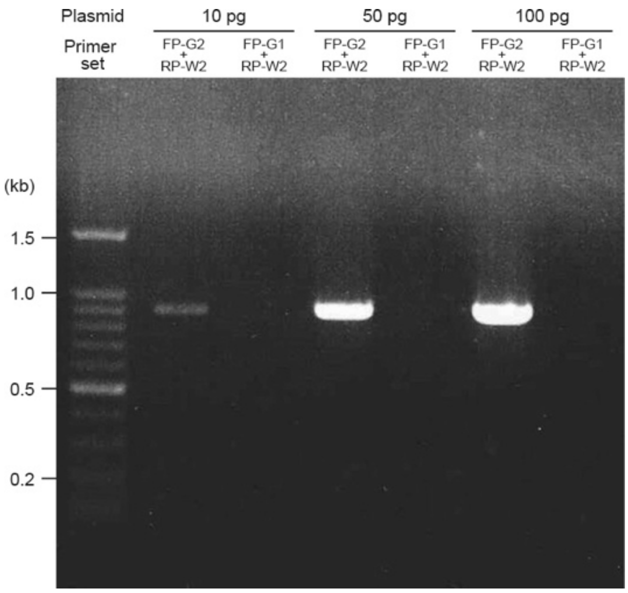


**Figure S3. PCR analysis of the transfer plasmid pAAV-DIO-GFP**

PCR amplification was carried out with different amounts (10, 50, and 100 pg) of the transfer plasmid as a template by using the following primer sets, and PCR products were subjected to 1% agarose gel electrophoresis. The primer set FP-G2/RP-W2 (see Figure 1A) gave a DNA band of ~0.9 kb, which are derived from the original transfer plasmid (corresponding to fragment A in Figure 1A). The primer set FP-G1/RP-W2 (see Figure 1B) did not detect any DNA bands originating from recombinants with the reversal of the inverted GFP gene cassette.

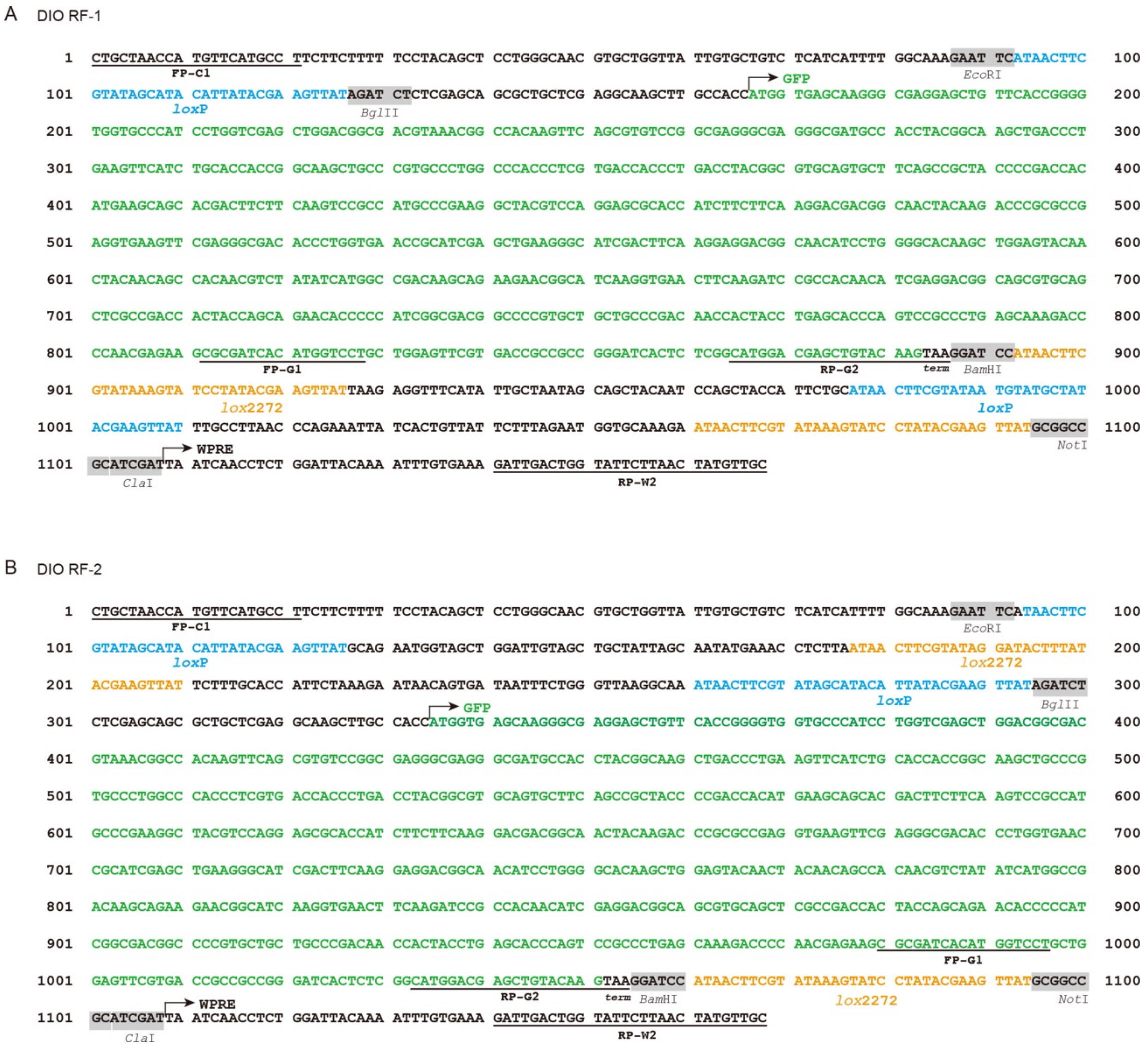


**Figure S4. Partial nucleotide sequence of viral genomes of RF-1 and RF-2 in AAV-DIO-GFP vector particles**

(A) Sequence of RF-1 genome obtained from the analysis of fragments B and E. (B) RF-2 sequence from the analysis of fragments C and F. Forward and reverse primer sequences are underlined.

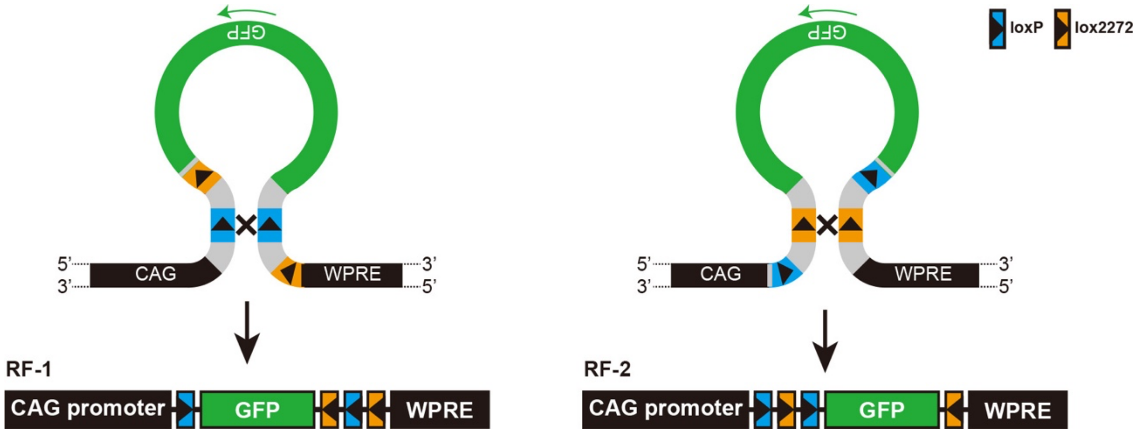


**Figure S5. Possible mechanisms that generate RF-1 and RF-2 genomes through recombination events during the production of AAV-DIO-GFP vector.**

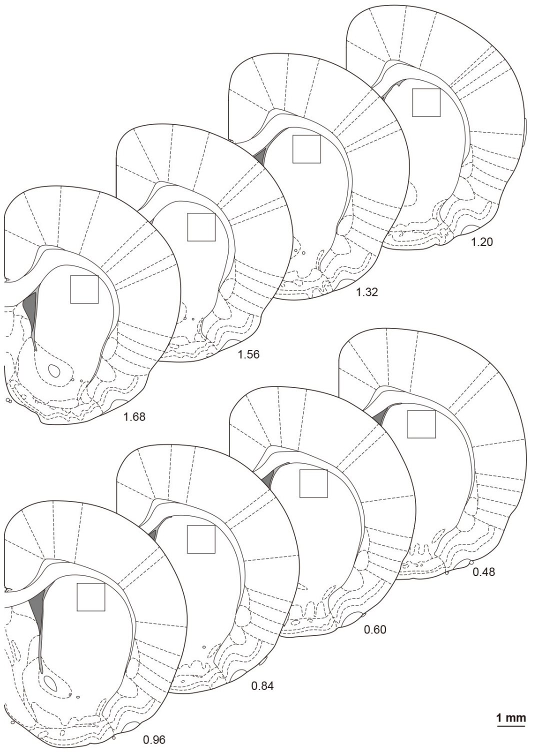


**Figure S6. Anteroposterior coordinates and ROIs used for cell counts**

Coronal sections through the rat striatum and ROIs (1.0 × 1.0 mm) are indicated with the coordinates (mm) from bregma. Scale bar: 1 mm.

A pAAV-DIO/5′USS-GFP

CAG promoter

1 GCGGCCGCAC GCGTCGACAT TGATTATTGA CTAGTTATTA ATAGTAATCA ATTACGGGGT CATTAGTTCA TAGCCCATAT ATGGAGTTCC GCGTTACATA 100

NotI PstI

101 ACTTACGGTA AATGGCCCGC CTGGCTGACC GCCCAACGAC CCCCGCCCAT TGACGTCAAT AATGACGTAT GTTCCCATAG TAACGCCAAT AGGGACTTTC 200

201 CATTGACGTC AATGGGTGGA GTATTTACGG TAAACTGCCC ACTTGGCAGT ACATCAAGTG TATCATATGC CAAGTACGCC CCCTATTGAC GTCAATGACG 300

301 GTAAATGGCC CGCCTGGCAT TATGCCCAGT ACATGACCTT ATGGGACTTT CCTACTTGGC AGTACATCTA CGTATTAGTC ATCGCTATTA CCATGGTCGA 400

401 GGTGAGCCCC ACGTTCTGCT TCACTCTCCC CATCTCCCCC CCCTCCCCAC CCCCAATTTT GTATTTATTT ATTTTTTAAT TATTTTGTGC AGCGATGGGG 500

501 GCGGGGGGGG GGGGGGGGCG CGCGCCAGGC GGGGCGGGGC GGGGCGAGGG GCGGGGCGGG GCGAGGCGGA GAGGTGCGGC GGCAGCCAAT CAGAGCGGCG 600

*

601 CGCTCCGAAA GTTTCCTTTT ATGGCGAGGC GGCGGCGGCG GCGGCCCTAT AAAAAGCGAA GCGCGCGGCG GGCGGGAGTC GCTGCGCGCT GCCTTCGCCC 700

701 CGTGCCCCGC TCCGCCGCCG CCTCGCGCCG CCCGCCCCGG CTCTGACTGA CCGCGTTACT CCCACAGgtg agcgggcggg acggcccttc tcctccgggc 800

801 tgtaattagc gcttggttta atgacggctt gtttctttc tgtggctgcg tgaaagcctt gaggggctcc gggagggccc tttgtgcggg gggagcggct 900

901 cgggggggtgc gtgcgtgtgt gtgtgcgtgg ggagcgccgc gtgcggctcc gcgctgcccg gcggctgtga gcgctgcggg cgcggcgcgg ggctttgtgc 1000

1001 gctccgcagt gtgcgcgagg ggagcgcggc cggggggcggt gccccgcggt gcggggggggg ctgcgagggg aacaaaggct gcgtgcgggg tgtgtgcgtg 1100

1101 gggggggtgag cagggggtgt gggcgcgtcg gtcgggctgc aaccccccct gcacccccct ccccgagttg ctgagcacgg cccggcttcg ggtgcggggc 1200

1201 tccgtacggg gcgtggcgcg gggctcgccg tgccgggcgg ggggtggcgg caggtggggg tgccgggcgg ggcggggccg cctcgggccg gggagggctc 1300

1301 gggggagggg cgcggcggcc cccggagcgc cggcggctgt cgaggcgcgg cgagccgcag ccattgcctt ttatggtaat cgtgcgagag ggcgcaggga 1400

1401 cttcctttgt cccaaatctg tgcggagccg aaatctggga ggcgccgccg caccccctct agcgggcgcg ggggcgaagcg gtgcggcgcc ggcaggaagg 1500

1501 aaatgggcgg ggagggcctt cgtgcgtcgc cgcgccgccg tccccttctc cctctccagc ctcggggctg tccgcggggg gacggctgcc ttcggggggg 1600

1601 acggggcagg gcggggttcg gcttctggcg tgtgaccggc ggctctagag cctctgctaa ccatgttcat gccttcttct ttttcctaca gCTCCTGGGC 1700

1701 AACGTGCTGG TTATTGTGCT GTCTCATCAT TTTGGCAAAG AATTCATAAC TTCGTATAGC ATACATTATA CGAAGTTATC CTAGCGCTAC CGGTCGCCAC 1800

EcoRI loxP 5′-USS AgeI

Turbo

1801 CATGGTGGGT GAGGATAGCG TGCTGATCAC CGAGAACATG CACATGAAAC TGTACATGGA GGGCACCGTG AACGACCACC ACTTCAAGTG CACATCCGAG 1900

1901 GGCGAAGGCA AGCCCTACGA GGGCACCCAG ACCATGAAGA TCAAGGTGGT CGAGGGCGGC CCTCTCCCCT TCGCCTTCGA CATCCTGGCT ACCAGCTTCA 2000

2001 TGTACGGCAG CAAAACCTTT ATCAACCACA CCCAGGGCAT CCCCGACTTC TTTAAGCAGT CCTTCCCTGA GGGCTTCACA TGGGAGAGGA TCACCACATA 2100

2101 CGAAGACGGG GGCGTGCTGA CCGCTACCCA GGACACCAGC CTCCAGAACG GCTGCCTCAT CTACAACGTC AAGATCAACG GGGTGAACTT CCCATCCAAC 2200

2201 GGCCCTGTGA TGCAGAAGAA AACACTCGGC TGGGAGGCCA GCACCGAGAT GCTGTACCCC GCTGACAGCG GCCTGAGAGG CCATAGCCAG ATGGCCCTGA 2300

2301 AGCTCGTGGG CGGGGGCTAC CTGCACTGCT CCCTCAAGAC CACATACAGA TCCAAGAAAC CCGCTAAGAA CCTCAAGATG CCCGGCTTCT ACTTCGTGGA 2400

2401 CAGGAGACTG GAAAGAATCA AGGAGGCCGA CAAAGAGACC TACGTCGAGC AGCACGAGAT GGCTGTGGCC AGGTACTGCG ACCTGCCTAG CAAACTGGGG 2500

term

2501 CACAGCTGAT GCGGCCGCGA CGCTAGCGCA TAACTTCGTA TAGGATACTT TATACGAAGT TATGGATCCT TACTTGTACA GCTCGTCCAT GCCGAGAGTG 2600

term NheI lox2272 BamHI

2601 ATCCCGGCGG CGGTCACGAA CTCCAGCAGG ACCATGTGAT CGCGCTTCTC GTTGGGGTCT TTGCTCAGGG CGGACTGGGT GCTCAGGTAG TGGTTGTCGG 2700

2701 GCAGCAGCAC GGGGCCGTCG CCGATGGGGG TGTTCTGCTG GTAGTGGTCG GCGAGCTGCA CGCTGCCGTC CTCGATGTTG TGGCGGATCT TGAAGTTCAC 2800

2801 CTTGATGCCG TTCTTCTGCT TGTCGGCCAT GATATAGACG TTGTGGCTGT TGTAGTTGTA CTCCAGCTTG TGCCCCAGGA TGTTGCCGTC CTCCTTGAAG 2900

2901 TCGATGCCCT TCAGCTCGAT GCGGTTCACC AGGGTGTCGC CCTCGAACTT CACCTCGGCG CGGGTCTTGT AGTTGCCGTC GTCCTTGAAG AAGATGGTGC 3000

3001 GCTCCTGGAC GTAGCCTTCG GGCATGGCGG ACTTGAAGAA GTCGTGCTGC TTCATGTGGT CGGGGTAGCG GCTGAAGCAC TGCACGCCGT AGGTCAGGGT 3100

3101 GGTCACGAGG GTGGGCCAGG GCACGGGCAG CTTGCCGGTG GTGCAGATGA ACTTCAGGGT CAGCTTGCCG TAGGTGGCAT CGCCCTCGCC CTCGCCGGAC 3200

3201 ACGCTGAACT TGTGGCCGTT TACGTCGCCG TCCAGCTCGA CCAGGATGGG CACCACCCCG GTGAACAGCT CCTCGCCCTT GCTCACCATG GTGGCAAGCT 3300

GFP

3301 TGCCTCGAGC AGCGCTGCTC GAGAGATCTA TAACTTCGTA TAATGTATGC TATACGAAGT TATTTGCCTT AACCCAGAAA TTATCACTGT TATTCTTTAG 3400

BglII loxP

WPRE

3401 AATGGTGCAA AGAATAACTT CGTATAAAGT ATCCTATACG AAGTTATGCG GCCGCATCGA Ttaatcaacc tctggattac aaaatttgtg aaagattgac 3500

lox2272 NotI ClaI

3501 tggtattctt aactatgttg ctccttttac gctatgtgga tacgctgctt taatgccttt gtatcatgct attgcttccc gtatggcttt cattttctcc 3600

3601 tccttgtata aatcctggtt gctgtctctt tatgaggagt tgtggcccgt tgtcaggcaa cgtggcgtgg tgtgcactgt gtttgctgac gcaaccccca 3700

3701 ctggttgggg cattgccacc acctgtcagc tcctttccgg gactttcgct ttccccctcc ctattgccac ggcggaactc atcgccgcct gccttgcccg 3800

3801 ctgctggaca ggggctcggc tgttgggcac tgacaattcc gtggtgttgt cggggaagct gacgtccttt ccatggctgc tcgcctgtgt tgccacctgg 3900

3901 attctgcgcg ggacgtcctt ctgctacgtc ccttcggccc tcaatccagc ggaccttcct tcccgcggcc tgctgccggc tctgcggcct cttccgcgtc 4000

hGH gene

4001 ttcgccttcg ccctcagacg agtcggatct ccctttgggc cgcctccccg catctgatct acgggtggca tccctgtgac ccctccccag tgcctctcct 4100

4101 ggccctggaa gttgccactc cagtgcccac cagccttgtc ctaataaaat taagttgcat cattttgtct gactaggtgt ccttctataa tattatgggg 4200

pA

4201 tggagggggg tggtatggag caaggggcaa gttgggaaga caacctgtag ggcctgcggg gtctattggg aaccaagctg gagtgcagtg gcacaatctt 4300

4301 ggctcactgc aatctccgcc tcctgggttc aagcgattct cctgcctcag cctcccgagt tgttgggatt ccaggcatgc atgaccaggc tcagctaatt 4400

4401 tttgtttttt tggtagagac ggggtttcac catattggcc aggctggtct ccaactccta atctcaggtg atctacccac cttggcctcc caaattgctg 4500

4501 ggattacagg cgtgaaccac tgctcccttc cctgtcctt

B pAAV-DIO/3'USS-GFP

```
        → CAG promoter
1    GCGGCCGCAC GCGTCGACAT TGATTATTGA CTAGTTATTA ATAGTAATCA ATTACGGGGT CATTAGTTCA TAGCCCATAT ATGGAGTTCC GCGTTACATA  100
      NotI       PstI
101  ACTTACGGTA AATGGCCCGC CTGGCTGACC GCCCAACGAC CCCCGCCCAT TGACGTCAAT AATGACGTAT GTTCCCATAG TAACGCCAAT AGGGACTTTC  200
201  CATTGACGTC AATGGGTGGA GTATTTACGG TAAACTGCCC ACTTGGCAGT ACATCAAGTG TATCATATGC CAAGTACGCC CCCTATTGAC GTCAATGACG  300
301  GTAAATGGCC CGCCTGGCAT TATGCCCAGT ACATGACCTT ATGGGACTTT CCTACTTGGC AGTACATCTA CGTATTAGTC ATCGCTATTA CCATGGTCGA  400
401  GGTGAGCCCC ACGTTCTGCT TCACTCTCCC CATCTCCCCC CCCTCCCCAC CCCCAATTTT GTATTTATTT ATTTTTTAAT TATTTTGTGC AGCGATGGGG  500
501  GCGGGGGGGG GGGGGGGGCG CGCGCCAGGC GGGGCGGGGC GGGGCGAGGG GCGGGGCGGG GCGAGGCGGA GAGGTGCGGC GGCAGCCAAT CAGAGCGGCG  600
                                                                                  *
601  CGCTCCGAAA GTTTCCTTTT ATGGCGAGGC GGCGGCGGCG GCGGCCCTAT AAAAAGCGAA GCGCGCGGCG GGCGGGAGTC GCTGCGCGCT GCCTTCGCCC  700
701  CGTGCCCCGC TCCGCCGCCG CCTCGCGCCG CCCGCCCCGG CTCTGACTGA CCGCGTTACT CCCACAGgtg agcgggcggg acggcccttc tcctccgggc  800
801  tgtaattagc gcttggttta atgacggctt gtttcttttc tgtggctgcg tgaaagcctt gaggggctcc gggagggccc tttgtgcggg gggagcggct  900
901  cgggggggtgc gtgcgtgtgt gtgtgcgtgg ggagcgccgc gtgcggctcc gcgctgcccg gcggctgtga gcgctgcggg cgcggcgcgg ggctttgtgc  1000
1001 gctccgcagt gtgcgcgagg ggagcgcggc cgggggcggt gccccgcggt gcgggggggg ctgcgagggg aacaaaggct gcgtgcgggg tgtgtgcgtg  1100
1101 ggggggtgag cagggggtgt gggcgcgtcg gtcgggctgc aacccccct gcacccccct ccccgagttg ctgagcacgg cccggcttcg ggtgcggggc  1200
1201 tccgtacggg gcgtggcgcg gggctcgccg tgccgggcgg ggggtggcgg caggtggggg tgccgggcgg ggcggggccg cctcgggccg gggagggctc  1300
1301 gggggagggg cgcggcggcc cccggagcgc cggcggctgt cgaggcgcgg cgagccgcag ccattgcctt ttatggtaat cgtgcgagag ggcgcaggga  1400
1401 cttcctttgt cccaaatctg tgcggagccg aaatctggga ggcgccgccg caccccctct agcgggcgcg gggcgaagcg gtgcggcgcc ggcaggaagg  1500
1501 aaatgggcgg ggagggcctt cgtgcgtcgc cgcgccgccg tccccttctc cctctccagc ctcggggctg tccgcggggg gacggctgcc ttcgggggg  1600
1601 acggggcagg gcggggttcg gcttctggcg tgtgaccggc ggctctagag cctctgctaa ccatgttcat gccttcttct ttttcctaca gCTCCTGGGC  1700
1701 AACGTGCTGG TTATTGTGCT GTCTCATCAT TTTGGCAAAG AATTCATAAC TTCGTATAGC ATACATTATA CGAAGTTATG CAGAATGGTA GCTGGATTGT  1800
                                            EcoRI                   loxP
1801 AGCTGCTATT AGCAATATGA AACCTCTTAA TAACTTCGTA TAGGATACTT TATACGAAGT TATGGATCCT TACTTGTACA GCTCGTCCAT GCCGAGAGTG  1900
                                               lox2272            BamHI
1901 ATCCCGGCGG CGGTCACGAA CTCCAGCAGG ACCATGTGAT CGCGCTTCTC GTTGGGGTCT TTGCTCAGGG CGGACTGGGT GCTCAGGTAG TGGTTGTCGG  2000
2001 GCAGCAGCAC GGGGCCGTCG CCGATGGGGG TGTTCTGCTG GTAGTGGTCG GCGAGCTGCA CGCTGCCGTC CTCGATGTTG TGGCGGATCT TGAAGTTCAC  2100
2101 CTTGATGCCG TTCTTCTGCT TGTCGGCCAT GATATAGACG TTGTGGCTGT TGTAGTTGTA CTCCAGCTTG TGCCCCAGGA TGTTGCCGTC CTCCTTGAAG  2200
2201 TCGATGCCCT TCAGCTCGAT GCGGTTCACC AGGGTGTCGC CCTCGAACTT CACCTCGGCG CGGGTCTTGT AGTTGCCGTC GTCCTTGAAG AAGATGGTGC  2300
2301 GCTCCTGGAC GTAGCCTTCG GGCATGGCGG ACTTGAAGAA GTCGTGCTGC TTCATGTGGT CGGGGTAGCG GCTGAAGCAC TGCACGCCGT AGGTCAGGGT  2400
2401 GGTCACGAGG GTGGGCCAGG GCACGGGCAG CTTGCCGGTG GTGCAGATGA ACTTCAGGGT CAGCTTGCCG TAGGTGGCAT CGCCCTCGCC CTCGCCGGAC  2500
2501 ACGCTGAACT TGTGGCCGTT TACGTCGCCG TCCAGCTCGA CCAGGATGGG CACCACCCCG GTGAACAGCT CCTCGCCCTT GCTCACCATG GTGGCAAGCT  2600
                                                                                               GFP ←
                                                                                                 → Turbo
2601 TGCCTCGAGC AGCGCTGCTC GAGAGATCTA TAACTTCGTA TAATGTATGC TATACGAAGT TATCCTAGCG CTACCGGTCG CCACCATGGT GGGTGAGGAT  2700
                          BglII               loxP                 3'-USS   AgeI
2701 AGCGTGCTGA TCACCGAGAA CATGCACATG AAACTGTACA TGGAGGGCAC CGTGAACGAC CACCACTTCA AGTGCACATC CGAGGGCGAA GGCAAGCCCT  2800
2801 ACGAGGGCAC CCAGACCATG AAGATCAAGG TGGTCGAGGG CGGCCCTCTC CCCTTCGCCT TCGACATCCT GGCTACCAGC TTCATGTACG GCAGCAAAAC  2900
2901 CTTTATCAAC CACACCCAGG GCATCCCCGA CTTCTTTAAG CAGTCCTTCC CTGAGGGCTT CACATGGGAG AGGATCACCA CATACGAAGA CGGGGGCGTG  3000
3001 CTGACCGCTA CCCAGGACAC CAGCCTCCAG AACGGCTGCC TCATCTACAA CGTCAAGATC AACGGGGTGA ACTTCCCATC CAACGGCCCT GTGATGCAGA  3100
3101 AGAAAACACT CGGCTGGGAG GCCAGCACCG AGATGCTGTA CCCCGCTGAC AGCGGCCTGA GAGGCCATAG CCAGATGGCC CTGAAGCTCG TGGGCGGGGG  3200
3201 CTACCTGCAC TGCTCCCTCA AGACCACATA CAGATCCAAG AAACCCGCTA AGAACCTCAA GATGCCCGGC TTCTACTTCG TGGACAGGAG ACTGGAAAGA  3300
3301 ATCAAGGAGG CCGACAAAGA GACCTACGTC GAGCAGCACG AGATGGCTGT GGCCAGGTAC TGCGACCTGC CTAGCAAACT GGGGCACAGC TGATGCGGCC  3400
                                                                                                  term
                                                                          → WPRE
3401 GCGACGCTAG CGCATAACTT CGTATAAAGT ATCCTATACG AAGTTATGCG GCCGCATCGA Ttaatcaacc tctggattac aaaatttgtg aaagattgac  3500
          NheI                lox2272                    NcoI   ClaI
3501 tggtattctt aactatgttg ctccttttac gctatgtgga tacgctgctt taatgccttt gtatcatgct attgcttccc gtatggcttt cattttctcc  3600
3601 tccttgtata aatcctggtt gctgtctctt tatgaggagt tgtggcccgt tgtcaggcaa cgtggcgtgg tgtgcactgt gtttgctgac gcaaccccca  3700
3701 ctggttgggg cattgccacc acctgtcagc tcctttccgg gactttcgct ttccccctcc ctattgccac ggcggaactc atcgccgcct gccttgcccg  3800
3801 ctgctggaca ggggctcggc tgttgggcac tgacaattcc gtggtgttgt cggggaagct gacgtccttt ccatggctgc tcgcctgtgt tgccacctgg  3900
3901 attctgcgcg ggacgtcctt ctgctacgtc ccttcggccc tcaatccagc ggaccttcct tcccgcggcc tgctgccggc tctgcggcct cttccgcgtc  4000
                                                                    → hGH gene
4001 ttcgccttcg ccctcagacg agtcggatct ccctttgggc cgcctccccg catctgatct acgggtggca tccctgtgac ccctccccag tgcctctcct  4100
4101 ggccctggaa gttgccactc cagtgcccac cagccttgtc ctaataaaat taagttgcat cattttgtct gactaggtgt ccttctataa tattatgggg  4200
                                                       pA
4201 tggagggggg tggtatggag caaggggcaa gttgggaaga caacctgtag ggcctgcggg gtctattggg aaccaagctg gagtgcagtg gcacaatctt  4300
4301 ggctcactgc aatctccgcc tcctgggttc aagcgattct cctgcctcag cctcccgagt tgttgggatt ccaggcatgc atgaccaggc tcagctaatt  4400
4401 tttgtttttt tggtagagac ggggtttcac catattggcc aggctggtct ccaactccta atctcaggtg atctacccac cttggcctcc caaattgctg  4500
4501 ggattacagg cgtgaaccac tgctcccttc cctgtcctt
```

**Figure S7. Nucleotide sequence of pAAV-DIO/5'USS-GFP and pAAV-DIO/3'USS-GFP transfer plasmids**

(A) Sequence of AAV-DIO/5'USS-GFP plasmid. (B) Sequence of AAV-DIO/3'USS-GFP plasmid. Asterisk indicates the transcription initiation site, and arrowheads show the splice donor and acceptor sites. The inverted GFP coding sequence, TurboFP635 coding sequence, *lox*P site and *lox*2272 site are shown by green, magenta, blue, and orange, respectively. 5'-USS and 3'-USS sequences are underlined. pA: hGH polyadenylation signal.

**A** DIO/5'USS RF-1

```
   1  CTGCTAACCA TGTTCATGCC TTCTTCTTTT TCCTACAGCT CCTGGGCAAC GTGCTGGTTA TTGTGCTGTC TCATCATTTT GGCAAAGAAT TCATAACTTC   100
           FP-C1                                                                                EcoRI
                                                                     GFP
 101  GTATAGCATA CATTATACGA AGTTATAGAT CTCTCGAGCA GCGCTGCTCG AGGCAAGCTT GCCACCATGG TGAGCAAGGG CGAGGAGCTG TTCACCGGGG   200
          loxP                  BglII
 201  TGGTGCCCAT CCTGGTCGAG CTGGACGGCG ACGTAAACGG CCACAAGTTC AGCGTGTCCG GCGAGGGCGA GGGCGATGCC ACCTACGGCA AGCTGACCCT   300
 301  GAAGTTCATC TGCACCACCG GCAAGCTGCC CGTGCCCTGG CCCACCCTCG TGACCACCCT GACCTACGGC GTGCAGTGCT TCAGCCGCTA CCCCGACCAC   400
 401  ATGAAGCAGC ACGACTTCTT CAAGTCCGCC ATGCCCGAAG GCTACGTCCA GGAGCGCACC ATCTTCTTCA AGGACGACGG CAACTACAAG ACCCGCGCCG   500
 501  AGGTGAAGTT CGAGGGCGAC ACCCTGGTGA ACCGCATCGA GCTGAAGGGC ATCGACTTCA AGGAGGACGG CAACATCCTG GGGCACAAGC TGGAGTACAA   600
 601  CTACAACAGC CACAACGTCT ATATCATGGC CGACAAGCAG AAGAACGGCA TCAAGGTGAA CTTCAAGATC CGCCACAACA TCGAGGACGG CAGCGTGCAG   700
 701  CTCGCCGACC ACTACCAGCA GAACACCCCC ATCGGCGACG GCCCCGTGCT GCTGCCCGAC AACCACTACC TGAGCACCCA GTCCGCCCTG AGCAAAGACC   800
 801  CCAACGAGAA GCGCGATCAC ATGGTCCTGC TGGAGTTCGT GACCGCCGCC GGGATCACTC TCGGCATGGA CGAGCTGTAC AAGTAAGGAT CCATAACTTC   900
                   FP-G1                                                   RP-G2       term  BamHI
                                                   term
 901  GTATAAAGTA TCCTATACGA AGTTATGCGC TAGCGTCGCG GCCGCATCAG CTGTGCCCCA GTTTGCTAGG CAGGTCGCAG TACCTGGCCA CAGCCATCTC  1000
          lox2272              NheI
1001  GTGCTGCTCG ACGTAGGTCT CTTTGTCGGC CTCCTTGATT CTTTCCAGTC TCCTGTCCAC GAAGTAGAAG CCGGGCATCT TGAGGTTCTT AGCGGGTTTC  1100
1101  TTGGATCTGT ATGTGGTCTT GAGGGAGCAG TGCAGGTAGC CCCCGCCCAC GAGCTTCAGG GCCATCTGGC TATGGCCTCT CAGGCCGCTG TCAGCGGGGT  1200
1201  ACAGCATCTC GGTGCTGGCC TCCCAGCCGA GTGTTTTCTT CTGCATCACA GGGCCGTTGG ATGGGAAGTT CACCCCGTTG ATCTTGACGT TGTAGATGAG  1300
1301  GCAGCCGTTC TGGAGGCTGG TGTCCTGGGT AGCGGTCAGC ACGCCCCCGT CTTCGTATGT GGTGATCCTC TCCCATGTGA AGCCCTCAGG GAAGGACTGC  1400
1401  TTAAAGAAGT CGGGGATGCC CTGGGTGTGG TTGATAAAGG TTTTGCTGCC GTACATGAAG CTGGTAGCCA GGATGTCGAA GGCGAAGGGG AGAGGGCCGC  1500
1501  CCTCGACCAC CTTGATCTTC ATGGTCTGGG TGCCCTCGTA GGGCTTGCCT TCGCCCTCGG ATGTGCACTT GAAGTGGTGG TCGTTCACGG TGCCCTCCAT  1600
1601  GTACAGTTTC ATGTGCATGT TCTCGGTGAT CAGCACGCTA TCCTCACCCA CCATGGTGGC GACCGGTAGC GCTAGGATAA CTTCGTATAA TGTATGCTAT  1700
                                                      Turbo          AgeI                         loxP
              WPRE
1701  ACGAAGTTAT TTGCCTTAAC CCAGAAATTA TCACTGTTAT TCTTTAGAAT GGTGCAAAGA ATAACTTCGT ATAAAGTATC CTATACGAAG TTATGCGGCC  1800
                                                                              lox2272               NotI
1801  GCATCGATTA ATCAACCTCT GGATTACAAA ATTTGTGAAA GATTGACTGG TATTCTTAAC TATGTTGC
        ClaI                                           RP-W2
```

**B** DIO/5'USS RF-2

```
   1  CTGCTAACCA TGTTCATGCC TTCTTCTTTT TCCTACAGCT CCTGGGCAAC GTGCTGGTTA TTGTGCTGTC TCATCATTTT GGCAAAGAAT TCATAACTTC   100
           FP-C1                                                                                EcoRI
                                                     Turbo
 101  GTATAGCATA CATTATACGA AGTTATCCTA GCGCTACCGG TCGCCACCAT GGTGGGTGAG GATAGCGTGC TGATCACCGA GAACATGCAC ATGAAACTGT   200
          loxP                             AgeI
 201  ACATGGAGGG CACCGTGAAC GACCACCACT TCAAGTGCAC ATCCGAGGGC GAAGGCAAGC CCTACGAGGG CACCCAGACC ATGAAGATCA AGGTGGTCGA   300
 300  GGGCGGCCCT CTCCCCTTCG CCTTCGACAT CCTGGCTACC AGCTTCATGT ACGGCAGCAA AACCTTTATC AACCACACCC AGGGCATCCC CGACTTCTTT   400
 401  AAGCAGTCCT TCCCTGAGGG CTTCACATGG GAGAGGATCA CCACATACGA AGACGGGGGC GTGCTGACCG CTACCCAGGA CACCAGCCTC CAGAACGGCT   500
 501  GCCTCATCTA CAACGTCAAG ATCAACGGGG TGAACTTCCC ATCCAACGGC CCTGTGATGC AGAAGAAAAC ACTCGGCTGG GAGGCCAGCA CCGAGATGCT   600
 601  GTACCCCGCT GACAGCGGCC TGAGAGGCCA TAGCCAGATG GCCCTGAAGC TCGTGGGCGG GGGCTACCTG CACTGCTCCC TCAAGACCAC ATACAGATCC   700
 701  AAGAAACCCG CTAAGAACCT CAAGATGCCC GGCTTCTACT TCGTGGACAG GAGACTGGAA AGAATCAAGG AGGCCGACAA AGAGACCTAC GTCGAGCAGC   800
 801  ACGAGATGGC TGTGGCCAGG TACTGCGACC TGCCTAGCAA ACTGGGGCAC AGCTGATGCG GCCGCGACGC TAGCGCATAA CTTCGTATAG GATACTTTAT   900
                                                               term           NheI                    lox2272
 901  ACGAAGTTAT TCTTTGCACC ATTCTAAAGA ATAACAGTGA TAATTTCTGG GTTAAGGCAA ATAACTTCGT ATAGCATACA TTATACGAAG TTATAGATCT  1000
                                     GFP                                             loxP                BglII
1001  CTCGAGCAGC GCTGCTCGAG GCAAGCTTGC CACCATGGTG AGCAAGGGCG AGGAGCTGTT CACCGGGGTG GTGCCCATCC TGGTCGAGCT GGACGGCGAC  1100
1101  GTAAACGGCC ACAAGTTCAG CGTGTCCGGC GAGGGCGAGG GCGATGCCAC CTACGGCAAG CTGACCCTGA AGTTCATCTG CACCACCGGC AAGCTGCCCG  1200
1201  TGCCCTGGCC CACCCTCGTG ACCACCCTGA CCTACGGCGT GCAGTGCTTC AGCCGCTACC CCGACCACAT GAAGCAGCAC GACTTCTTCA AGTCCGCCAT  1300
1301  GCCCGAAGGC TACGTCCAGG AGCGCACCAT CTTCTTCAAG GACGACGGCA ACTACAAGAC CCGCGCCGAG GTGAAGTTCG AGGGCGACAC CCTGGTGAAC  1400
1401  CGCATCGAGC TGAAGGGCAT CGACTTCAAG GAGGACGGCA ACATCCTGGG GCACAAGCTG GAGTACAACT ACAACAGCCA CAACGTCTAT ATCATGGCCG  1500
1501  ACAAGCAGAA GAACGGCATC AAGGTGAACT TCAAGATCCG CCACAACATC GAGGACGGCA GCGTGCAGCT CGCCGACCAC TACCAGCAGA ACACCCCCAT  1600
1601  CGGCGACGGC CCCGTGCTGC TGCCCGACAA CCACTACCTG AGCACCCAGT CCGCCCTGAG CAAAGACCCC AACGAGAAGC GCGATCACAT GGTCCTGCTG  1700
                                                                                                FP-G1
1701  GAGTTCGTGA CCGCCGCCGG GATCACTCTC GGCATGGACG AGCTGTACAA GTAAGGATCC ATAACTTCGT ATAAAGTATC CTATACGAAG TTATGCGGCC  1800
              WPRE                                    RP-G2       term  BamHI          lox2272               NotI
1801  GCATCGATTA ATCAACCTCT GGATTACAAA ATTTGTGAAA GATTGACTGG TATTCTTAAC TATGTTGC
        ClaI                                           RP-W2
```

**C** DIO/3'USS RF-1

```
1     CTGCTAACCA TGTTCATGCC TTCTTCTTTT TCCTACAGCT CCTGGGCAAC GTGCTGGTTA TTGTGCTGTC TCATCATTTTG GCAAAGAAT TCATAACTTC   100
          FP-C1                                                                                   GFP        EcoRI
101   GTATAGCATA CATTATACGA AGTTATAGAT CTCTCGAGCA GCGCTGCTCG AGGCAAGCTT GCCACCATGG TGAGCAAGGG CGAGGAGCTG TTCACCGGGG   200
          loxP             BglII
201   TGGTGCCCAT CCTGGTCGAG CTGGACGGCG ACGTAAACGG CCACAAGTTC AGCGTGTCCG GCGAGGGCGA GGGCGATGCC ACCTACGGCA AGCTGACCCT   300
301   GAAGTTCATC TGCACCACCG GCAAGCTGCC CGTGCCCTGG CCCACCCTCG TGACCACCCT GACCTACGGC GTGCAGTGCT TCAGCCGCTA CCCCGACCAC   400
401   ATGAAGCAGC ACGACTTCTT CAAGTCCGCC ATGCCCGAAG GCTACGTCCA GGAGCGCACC ATCTTCTTCA AGGACGACGG CAACTACAAG ACCCGCGCCG   500
501   AGGTGAAGTT CGAGGGCGAC ACCCTGGTGA ACCGCATCGA GCTGAAGGGC ATCGACTTCA AGGAGGACGG CAACATCCTG GGGCACAAGC TGGAGTACAA   600
601   CTACAACAGC CACAACGTCT ATATCATGGC CGACAAGCAG AAGAACGGCA TCAAGGTGAA CTTCAAGATC CGCCACAACA TCGAGGACGG CAGCGTGCAG   700
701   CTCGCCGACC ACTACCAGCA GAACACCCCC ATCGGCGACG GCCCCGTGCT GCTGCCCGAC AACCACTACC TGAGCACCCA GTCCGCCCTG AGCAAAGACC   800
801   CCAACGAGAA GCGCGATCAC ATGGTCCTGC TGGAGTTCGT GACCGCCGCC GGGATCACTC TCGGCATGGA CGAGCTGTAC AAGTAAGGAT CCATAACTTC   900
                    FP-G1                                             RP-G2         term BamHI
901   GTATAAAGTA TCCTATACGA AGTTATTAAG AGGTTTCATA TTGCTAATAG CAGCTACAAT CCAGCTACCA TTCTGCATAA CTTCGTATAA TGTATGCTAT  1000
          lox2272                                                                                     loxP
1001  ACGAAGTTAT CCTAGCGCTA CCGGTCGCCA CCATGGTGGG TGAGGATAGC GTGCTGATCA CCGAGAACAT GCACATGAAA CTGTACATGG AGGGCACCGT  1100
                     AgeI          Turbo
1101  GAACGACCAC CACTTCAAGT GCACATCCGA GGGCGAAGGC AAGCCCTACG AGGGCACCCA GACCATGAAG ATCAAGGTGG TCGAGGGCGG CCCTCTCCCC  1200
1201  TTCGCCTTCG ACATCCTGGC TACCAGCTTC ATGTACGGCA GCAAAACCTT TATCAACCAC ACCCAGGGCA TCCCCGACTT CTTTAAGCAG TCCTTCCCTG  1300
1301  AGGGCTTCAC ATGGGAGAGG ATCACCACAT ACGAAGACGG GGGCGTGCTG ACCGCTACCC AGGACACCAG CCTCCAGAAC GGCTGCCTCA TCTACAACGT  1400
1401  CAAGATCAAC GGGGTGAACT TCCCATCCAA CGGCCCTGTG ATGCAGAAGA AAACACTCGG CTGGGAGGCC AGCACCGAGA TGCTGTACCC CGCTGACAGC  1500
1501  GGCCTGAGAG GCCATAGCCA GATGGCCCTG AAGCTCGTGG GCGGGGGCTA CCTGCACTGC TCCCTCAAGA CCACATACAG ATCCAAGAAA CCCGCTAAGA  1600
1601  ACCTCAAGAT GCCCGGCTTC TACTTCGTGG ACAGGAGACT GGAAAGAATC AAGGAGGCCG ACAAAGAGAC CTACGTCGAG CAGCACGAGA TGGCTGTGGC  1700
1701  CAGGTACTGC GACCTGCCTA GCAAACTGGG GCACAGCTGA TGCGGCCGCG ACGCTAGCGC ATAACTTCGT ATAAAGTATC CTATACGAAG TTATGCGGCC  1800
                                          term              AgeI                lox2272               NotI
          WPRE
1801  GCATCGATTA ATCAACCTCT GGATTACAAA ATTTGTGAAA GATTGACTGG TATTCTTAAC TATGTTGC
        ClaI                                                RP-W2
```

**D** DIO/3'USS RF-2

```
1     CTGCTAACCA TGTTCATGCC TTCTTCTTTT TCCTACAGCT CCTGGGCAAC GTGCTGGTTA TTGTGCTGTC TCATCATTTT GGCAAAGAAT TCATAACTTC   100
          FP-C1                                                                                              EcoRI
101   GTATAGCATA CATTATACGA AGTTATGCAG AATGGTAGCT GGATTGTAGC TGCTATTAGC AATATGAAAC CTCTTAATAA CTTCGTATAG GATACTTTAT   200
          loxP                        term                                                         lox2272
201   ACGAAGTTAT GCGCTAGCGT CGCGGCCGCA TCAGCTGTGC CCCAGTTTGC TAGGCAGGTC GCAGTACCTG GCCACAGCCA TCTCGTGCTG CTCGACGTAG   300
                   NheI
301   GTCTCTTTGT CGGCCTCCTT GATTCTTTCC AGTCTCCTGT CCACGAAGTA GAAGCCGGGC ATCTTGAGGT TCTTAGCGGG TTTCTTGGAT CTGTATGTGG   400
401   TCTTGAGGGA GCAGTGCAGG TAGCCCCCGC CCACGAGCTT CAGGGCCATC TGGCTATGGC CTCTCAGGCC GCTGTCAGCG GGGTACAGCA TCTCGGTGCT   500
501   GGCCTCCCAG CCGAGTGTTT TCTTCTGCAT CACAGGGCCG TTGGATGGGA AGTTCACCCC GTTGATCTTG ACGTTGTAGA TGAGGCAGCC GTTCTGGAGG   600
601   CTGGTGTCCT GGGTAGCGGT CAGCACGCCC CCGTCTTCGT ATGTGGTGAT CCTCTCCCAT GTGAAGCCCT CAGGGAAGGA CTGCTTAAAG AAGTCGGGGA   700
701   TGCCCTGGGT GTGGTTGATA AAGGTTTTGC TGCCGTACAT GAAGCTGGTA GCCAGGATGT CGAAGGCGAA GGGGAGAGGG CCGCCCTCGA CCACCTTGAT   800
801   CTTCATGGTC TGGGTGCCCT CGTAGGGCTT GCCTTCGCCC TCGGATGTGC ACTTGAAGTG GTGGTCGTTC ACGGTGCCCT CCATGTACAG TTTCATGTGC   900
901   ATGTTCTCGG TGATCAGCAC GCTATCCTCA CCCACCATGG TGGCGACCGG TAGCGCTAGG ATAACTTCGT ATAGCATACA TTATACGAAG TTATAGATCT  1000
                                       Turbo          AgeI                    loxP                  BglII
                                             GFP
1001  CTCGAGCAGC GCTGCTCGAG GCAAGCTTGC CACCATGGTG AGCAAGGGCG AGGAGCTGTT CACCGGGGTG GTGCCCATCC TGGTCGAGCT GGACGGCGAC  1100
1101  GTAAACGGCC ACAAGTTCAG CGTGTCCGGC GAGGGCGAGG GCGATGCCAC CTACGGCAAG CTGACCCTGA AGTTCATCTG CACCACCGGC AAGCTGCCCG  1200
1201  TGCCCTGGCC CACCCTCGTG ACCACCCTGA CCTACGGCGT GCAGTGCTTC AGCCGCTACC CCGACCACAT GAAGCAGCAC GACTTCTTCA AGTCCGCCAT  1300
1301  GCCCGAAGGC TACGTCCAGG AGCGCACCAT CTTCTTCAAG GACGACGGCA ACTACAAGAC CCGCGCCGAG GTGAAGTTCG AGGGCGACAC CCTGGTGAAC  1400
1401  CGCATCGAGC TGAAGGGCAT CGACTTCAAG GAGGACGGCA ACATCCTGGG GCACAAGCTG GAGTACAACT ACAACAGCCA CAACGTCTAT ATCATGGCCG  1500
1501  ACAAGCAGAA GAACGGCATC AAGGTGAACT TCAAGATCCG CCACAACATC GAGGACGGCA GCGTGCAGCT CGCCGACCAC TACCAGCAGA ACACCCCCAT  1600
1601  CGGCGACGGC CCCGTGCTGC TGCCCGACAA CCACTACCTG AGCACCCAGT CCGCCCTGAG CAAAGACCCC AACGAGAAGC GCGATCACAT GGTCCTGCTG  1700
                                                                                                   FP-G1
1701  GAGTTCGTGA CCGCCGCCGG GATCACTCTC GGCATGGACG AGCTGTACAA GTAAGGATCC ATAACTTCGT ATAAAGTATC CTATACGAAG TTATGCGGCC  1800
                                                   RP-G2       term BamHI                   lox2272               NotI
          WPRE
1801  GCATCGATTA ATCAACCTCT GGATTACAAA ATTTGTGAAA GATTGACTGG TATTCTTAAC TATGTTGC
        ClaI                                                RP-W2
```

**Figure S8. Partial nucleotide sequence of RF-1/RF-2 genomes in AAV-DIO/5'USS-GFP and AAV- DIO/3'USS-GFP vector preparations.**

(A) Sequence of RF-1 genome obtained from the analysis of fragments B and H. (B) RF-2 sequence from the analysis of fragments C and I. (C) Sequence of RF-1 obtained from the analysis of fragments E and K. (D) RF-2 sequence from the analysis of fragments F and L. Forward and reverse primer sequences are underlined.

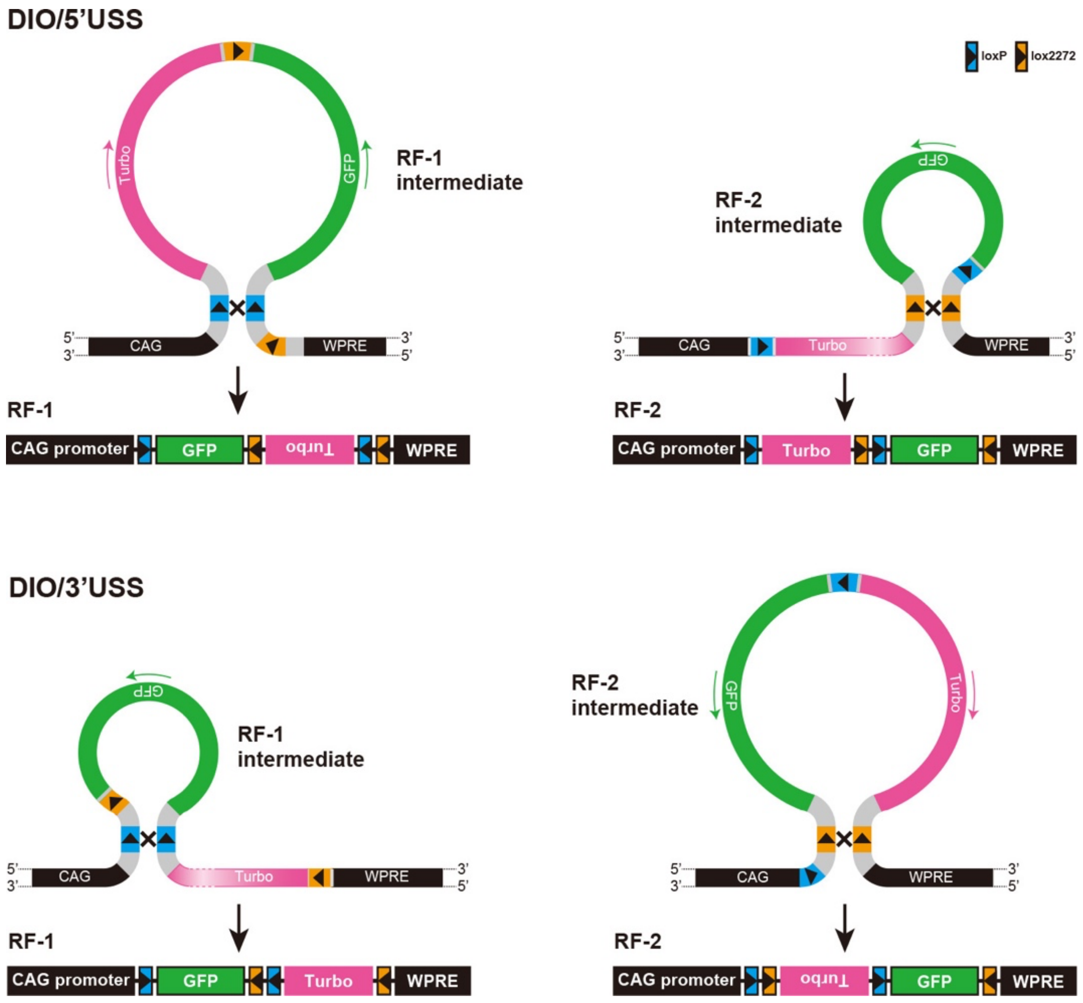


**Figure S9. Possible mechanisms that generate RF-1 and RF-2 genomes through recombination events during the production of AAV-DIO/5'USS-GFP and AAV-DIO/ 3'USS-GFP vectors.**

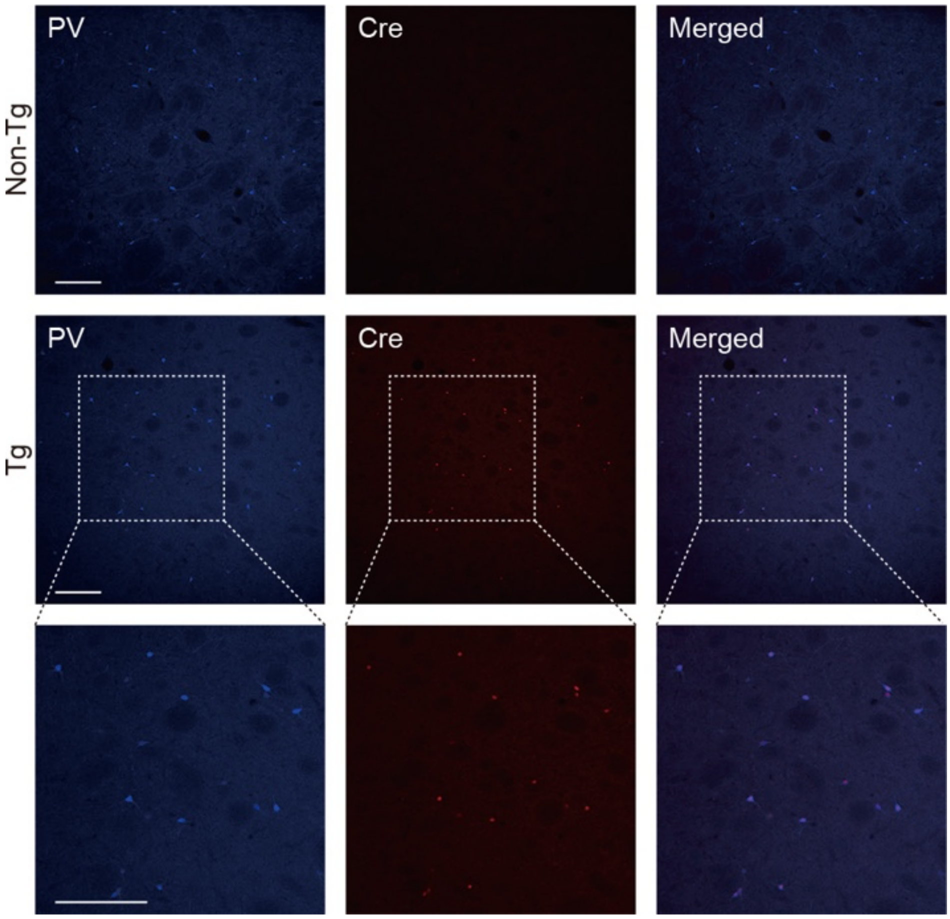


**Figure S10. Expression pattern of Cre transgene in the striatum of PV-Cre knock-in transgenic rats**

Sections through the striatum prepared from non-transgenic (non-Tg) and knock-in transgenic (Tg) rats were stained by double immunohistochemistry with anti-PV and anti-Cre antibodies. $Cre^+$ signals were visualized in all $PV^+$ cells in the knock-in transgenic rats. Scale bar: 200 μm.

```
                              → Turbo
    1 CCTAGCGCTA CCGGTCGCCA CCATGGTGGG TGAGGATAGC GTGCTGATCA CCGAGAACAT GCACATGAAA CTGTACATGG AGGGCACCGT GAACGACCAC  100
        Eco47III  AgeI        TAG                                    TA G   TAG          TAG
  101 CACTTCAAGT GCACATCCGA GGGCGAAGGC AAGCCCTACG AGGGCACCCA GACCATGAAG ATCAAGGTGG TCGAGGGCGG CCCTCTCCCC TTCGCCTTCG  200
              ApaLI                                              TAG
  201 ACATCCTGGC TACCAGCTTC ATGTACGGCA GCAAAACCTT TATCAACCAC ACCCAGGGCA TCCCCGACTT CTTTAAGCAG TCCTTCCCTG AGGGCTTCAC  300
                            TAG                                                                  Bsu36I
  301 ATGGGAGAGG ATCACCACAT ACGAAGACGG GGGCGTGCTG ACCGCTACCC AGGACACCAG CCTCCAGAAC GGCTGCCTCA TCTACAACGT CAAGATCAAC  400

  401 GGGGTGAACT TCCCATCCAA CGGCCCTGTG ATGCAGAAGA AAACACTCGG CTGGGAGGCC AGCACCGAGA TGCTGTACCC CGCTGACAGC GGCCTGAGAG  500
                                       TAG                                       T AG
  501 GCCATAGCCA GATGGCCCTG AAGCTCGTGG GCGGGGGCTA CCTGCACTGC TCCCTCAAGA CCACATACAG ATCCAAGAAA CCCGCTAAGA ACCTCAAGAT  600
                  TAG                                                                                             TA
  601 GCCCGGCTTC TACTTCGTGG ACAGGAGACT GGAAAGAATC AAGGAGGCCG ACAAAGAGAC CTACGTCGAG CAGCACGAGA TGGCTGTGGC CAGGTACTGC  700
      G                                                            PshAI                    T AG
  701 GACCTGCCTA GCAAACTGGG GCACAGCTGA TGCGGCCGCG ACGCTAGCGC
                                   term             NheI
```

**Figure S11. Nucleotide sequence of the TurboFP635 gene cassette**

Restriction enzyme sites used for the shortening of DNA fragment are shown. Eleven in-frame ATG codons, which are converted to the termination codon TAG, are underlined.

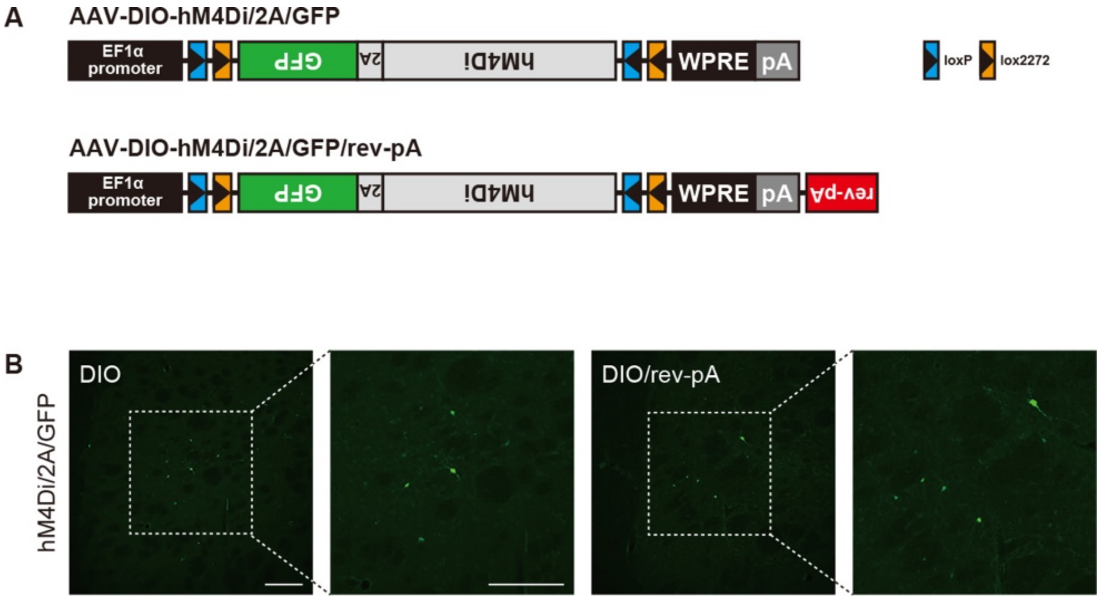


**Figure S12. Exclusion of the possibility of gene expression based on transcription from the inverted transgene**

AAV vector encoding a gene cassette containing rabbit $\beta$-globin gene polyadenylation signal in the reverse orientation (rev-pA) downstream of hGH gene polyadenylation signal (pA) of the transfer plasmid (termed AAV-DIO-hM4Di/2A/GFP/rev-pA) and AAV-DIO-hM4Di/2A/GFP vector as a control ($2.4 \times 10^{12}$ copies/mL) were used for intracranial injection into the rat striatum, and striatal sections were immunostained for GFP. (A) Genome structures of AAV-DIO-hM4Di/2A/GFP and AAV-DIO-hM4Di/2A/GFP/rev-pA vectors. (B) Representative photos of stained sections. Scale bars: 200 μm. The number of immuno-positive cells was similar between the two groups (4.95 ± 0.55 cells for AAV-DIO-hM4Di/2A/GFP and 5.02 ± 1.02 cells for AAV-DIO-hM4Di/2A/GFP/rev-pA, n = 4 animals), showing no significant difference between the groups (Student's $t$ test, $t_6 = 0.058$, $P = 0.955$).
